# SPINDLE: Unlocking protein dynamics from single-field NMR relaxation data using a deep learning ensemble

**DOI:** 10.64898/2026.07.31.742133

**Authors:** Olivia E. Krise, Michael P. Latham

## Abstract

A protein’s function is derived from its three-dimensional structure and the motions of the atoms about that structure. The detailed characterization of both macromolecular structure and dynamics provides an opportunity for understanding enzyme catalysis, ligand binding, and allostery, along with providing insights into how the function changes upon mutation or post-translational modification. Among the various methods for characterizing biomolecular motions, nuclear magnetic resonance (NMR) spin relaxation methods are a standard for determining nanosecond global tumbling times along with the amplitude and timescale of faster local motions. Within the model-free formalism, various mathematical models are used to extract dynamic parameters. Unfortunately, as the number of fitted parameters increases within these models, they become mathematically underdetermined for standard NMR relaxation data collected at a single magnetic field, necessitating multi-field datasets. Here, we present SPINDLE, an ensemble of deep neural networks trained on a large synthetic set of NMR relaxation data. Unlike traditional least-squares fitting, SPINDLE predicts both fast and slow timescale dynamics parameters from a single set (i.e., collected at a single magnetic field) of three relaxation datasets using the ensemble for error estimation. We demonstrate a strong correlation to ground truth dynamics parameters on synthetic benchmarks, with more precision than traditional fitting techniques, and precisely reproduce experimental dynamics parameters for ∼50 proteins with relaxation data in the Biological Magnetic Resonance Data Bank. We also leverage the architecture of the deep neural network to show how the model emphasizes rigid residues for the prediction of global correlation times. This strategy may be useful in the future for elucidating correlated networks of dynamic residues from multiple relaxation datasets.

## Introduction

The function of a protein results from the interplay of its three-dimensional structure and the motions that are superimposed upon that. Although several experimental (e.g., X-ray crystallography, cryogenic electron microscopy, and nuclear magnetic resonance [NMR] spectroscopy) and computational (e.g., AlphaFold^1^ and RoseTTaFold^2^) methods exist for determining the three-dimensional structure of biomolecules, relatively fewer methods are capable of site-specifically probing protein motions over the entire range of functionally relevant timescales. Among these methods for studying dynamics, NMR spectroscopy stands out for its ability to site-specifically probe functionally important biomolecular motions across timescales that range from picoseconds to days^3,4^. In fact, NMR spectroscopy has been instrumental in determining the importance of protein motion in enzyme catalysis^5,6^, ligand binding^7^, protein folding^8,9^, and allostery^10,11^, and these and similar studies have fundamentally changed the way we view macromolecular structure, dynamics, and function.

One of the most common and widely used frameworks for interpreting molecular motions from NMR data is through the analysis of spin relaxation measurements, particularly those utilizing the backbone amide ^15^N, using the Lipari-Szabo model-free formalism^12–14^. Here, relatively straightforward measurements of site-specific ^15^N R_1_ (longitudinal) and R_2_ (transverse) relaxation rates, describing the return of magnetization to equilibrium and the loss of magnetization in the x/y-plane, respectively, along with the {^1^H}-^15^N heteronuclear nuclear Overhauser effect (NOE) are mapped to various descriptors of molecular motion describing the global timescale of molecular tumbling (τ_c_) and local amplitude (S^2^ – the generalized order parameter) and timescales (τ_e_ and τ_s_ – fast and slow effective local correlation times, respectively) of fast ps-ns motions along with slow µs-ms timescale conformational exchange dynamics (R_ex_). Although the nomenclature can be confusing, the model-free formalism makes use of numerous mathematical models with different dynamics parameters to describe the experimental data (note, the mathematical models and their parameters are independent of the actual nature of the motions, hence model-free). For example, model 1 simply uses the motional parameters τ_c_ and S^2^ to describe the underlying motions responsible for the relaxation data, whereas model 2 (the original model proposed by Lipari and Szabo^12^) includes τ_c_, S^2^, and τ_e_, and model 4 (“extended” model-free^15^) can include τ_c_ and different order parameters and local correlation times for two timescales of motions.

The typical workflow for selecting the correct mathematical model for extracting dynamics parameters from experimental relaxation data relies on fitting the data to increasingly complex models and comparing the statistics of the fit (e.g., through F-test or AIC) to determine if the more complicated model is valid^16,17^. This process is repeated until there is no longer a statistical justification for increasing the complexity of the model. Several programs or wrappers exist to streamline the still time-consuming process for the user^16,18–20^. Regardless, a non-linear least squares fitting problem exists when analyzing these data: at one static magnetic field, there are typically three observables (i.e., ^15^N R_1_, ^15^N R_2_, and {^1^H}-^15^N NOE), which allows for accurate fitting of two unknowns. Thus, single-field data is appropriate for fitting residue-specific parameters from models 1 (S^2^), 2 (S^2^ and τ_e_), or 3 (S^2^ and R_ex_). Fitting to more complicated models, therefore, requires collecting data at additional static magnetic fields, a potentially time consuming and expensive endeavor that requires the sample to be stable for weeks. Moreover, extracting the conformational exchange term R_ex_ from spin dynamics data is also challenging even with two field relaxation data. Since R_ex_ adds to the intrinsic R_2_, it mathematically mimics the spectral density of a slower tumbling molecule or of a more rigid residue. As a result, traditional least squares-based optimizers get stuck in local minima when trying to fit this parameter.

Although the model-free approach is mathematically underdetermined when single-field relaxation data is used for extracting dynamics parameters from more complicated models (e.g., model 4: residue specific S^2^, τ_e_, and R_ex_ and global τ_c_), we hypothesized that machine learning algorithms, which can be trained on a vast number of easy to generate synthetic datasets representing the entire landscape of theoretical possibilities, can learn to separate the signals of the relaxation parameters globally. This training approach has been successfully applied to several applications for NMR spectroscopy including pulse design^21,22^, denoising^23^, automatic phasing^24,25^, peak picking^26^, processing non-uniformly sampled data^27–29^, and increasing spectral resolution from coupled spectra^30,31^. For example, Brüshweiler and co-workers trained DEEP picker against thousands of 1-dimensional spectra with varying degrees of peak overlap^26^. Interestingly, their models were readily expandable to peak picking multi-dimensional NMR data. In exciting applications combining the strengths of machine learning with clever NMR experiment implementation, Hansen, Vallupaluri, and co-workers have leveraged the ability to simulate vast quantities of NMR data to train machine learning algorithms to predict ‘normal-looking’ 2-dimensional ^13^C,^1^H NMR spectra from experimental data collected with only a single pulse^32^. Prior to the current wave of machine learning algorithms applications to NMR data processing, neural networks have been used to increase the accuracy of chemical shift^33^ and backbone torsion angle^34^ predictions, amino acid assignment,^35^ and peak picking^36,37^. Given the flexibility and eventual ease of use for the end user, it is surprising that machine learning approaches have not been more widely applied to downstream NMR data analysis applications, although notable applications include the peak picking, assignment, and structure calculation pipeline in ARTINA^38^ and the estimation of starting values for traditional non-linear least squares fitting of relaxation dispersion data in RING^39^.

Herein, we present SPINDLE (SPIN dynamics from Deep Learning Ensemble) – a python-based deep neural network (DNN) for predicting ^15^N dynamics parameters describing motions on the fast (ps-to-ns) and slow (µs-to-ms) timescales from standard ^15^N R_1_, ^15^N R_2_, and heteronuclear {^1^H}-^15^N NOE relaxation data collected at a single static magnetic field. Through training on a large volume of training data with synthetic ‘proteins’ varying in size from 40 to 450 ‘amino acids,’ we demonstrate that the DNN models contain the necessary information to accurately and quickly predict the global correlation time and three local dynamics parameters (S^2^, τ_e_, and R_ex_) for protein backbone amide ^15^N groups from only three experimental observables. To obtain errors in the predicted dynamics parameters, we used a bootstrap approach to generate multiple DNN models, and we utilize the variance across the DNN ensemble, which represents the epistemic uncertainty of the model, to provide robust error estimates for the predicted dynamics parameters, bypassing the need for the computationally expensive Monte Carlo simulations used by the standard fitting methods. The ensemble approach provides fast, robust error bounds for all the extracted dynamics parameters and allows for flagging ambiguous or contradictory predictions. Using additional synthetic datasets, we show that SPINDLE is faster and more accurate than standard least-squares fitting approaches, does not suffer from getting trapped in local minima, and foregoes the tedious and lengthy process of model selection. SPINDLE precisely captures τ_c_ for ∼50 proteins whose ^15^N relaxation data has been deposited into the Biological Magnetic Resonance Data Bank (BMRB). Moreover, the DNN provides excellent agreement with traditionally obtained S^2^ values and good agreement with R_ex_, whereas a weaker agreement with deposited τ_e_ values were observed. Finally, we show how a self-attention layer used in the DNN identifies rigid (i.e., high S^2^ and low R_ex_) residues within the protein sequence. Given the functionality and ease of use, SPINDLE should facilitate the analysis of NMR spin relaxation measurements for experts and non-experts alike.

## RESULTS AND DISCUSSION

### SPINDLE Architecture and Training

To predict both residue-specific local and global dynamics from variable-length NMR ^15^N spin relaxation data, we developed a multi-task deep learning architecture utilizing Bidirectional Long Short-Term Memory (LSTM) networks and a multi-head self-attention mechanism (Figure 1). The model accepts variable-length sequences of three experimental spin relaxation values (^15^N R_1_ and R_2_ rates and {^1^H}-^15^N heteronuclear NOE values) from a single static magnetic field and employs a masking layer to accommodate varying protein lengths without requiring fixed-size inputs. The core feature extraction block consists of two stacked Bidirectional LSTM layers. These layers process the relaxation data in both the forward and backward directions, allowing the network to capture contextual dynamics dependencies and long-range interactions along the protein sequence. This shared sequential representation is passed through a fully connected dense layer and normalized before splitting into two distinct predictive branches. The first branch maintains the sequence dimension, using a fully connected output layer to directly map the shared representation to the individual residues. Therefore, this first branch simultaneously predicts local generalized order parameter (S^2^), internal correlation time (τ_e_), and chemical exchange rate (R_ex_) for each residue in the sequence. To predict the isotropic global rotational correlation time (τ_c_), the shared sequence representation is passed through a multi-head self-attention layer. This attention mechanism allows the model to dynamically weight the most informative residues, effectively learning which parts of the protein are most indicative of overall tumbling. The attention-weighted sequence is condensed via 1D global average pooling and passed through a dropout-regularized dense layer to yield a single, global τ_c_ prediction for the entire protein.

**Figure 1.**
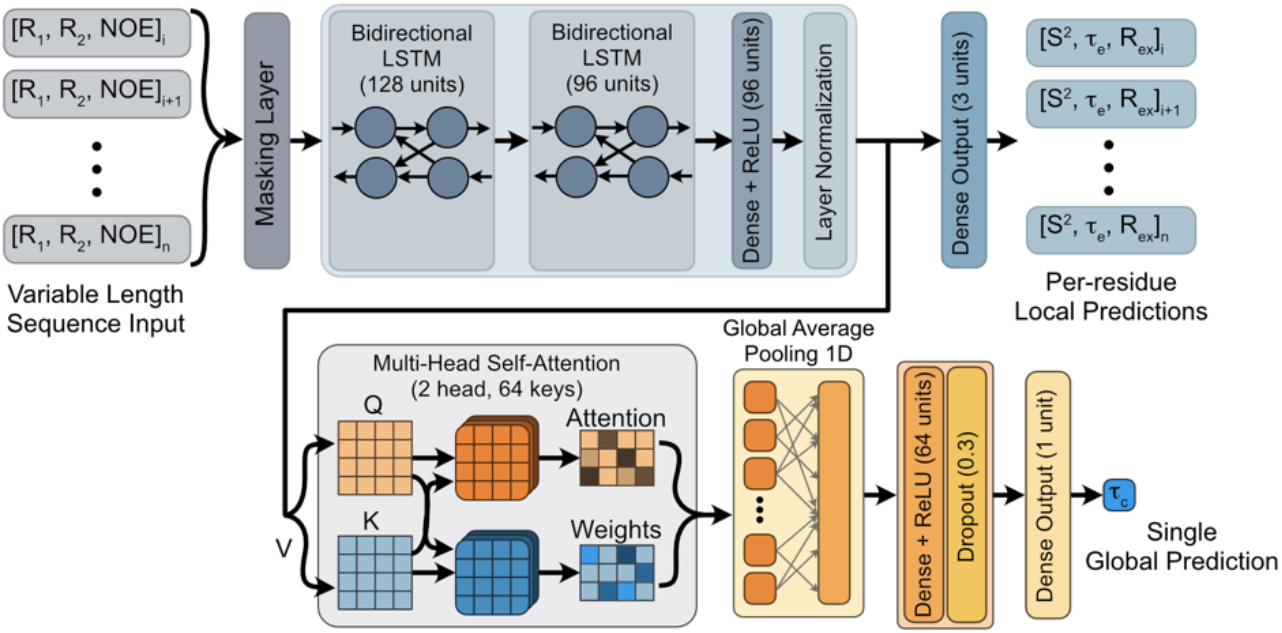
Architecture of the SPINDLE DNN for predicting NMR relaxation dynamics. The model takes a variable-length sequence of relaxation parameters (^15^N R_1_, R_2_, and NOE) as input. After masking, the input is processed through two Bidirectional LSTM layers, followed by a dense layer and a normalization step. The network then splits into two different prediction branches. The local branch applies a dense layer directly to the normalized sequence tensor to output per-residue predictions for the local dynamic parameters S^2^, τ_e_, and R_ex_. The global branch passes the same normalized sequence tensor into a Multi-Head Self-Attention layer to capture sequence-wide dependencies, where V is the value, Q is the query, and K is the key for the layer. The attended features are collapsed via 1D global average pooling, passed through dense and dropout layers, and finally processed by a dense layer to predict a single global rotational correlation time (τ_c_) for the entire sequence.

For different static magnetic field strengths between 500 MHz and 1100 MHz (^1^H Larmor frequencies), the deep neural network (DNN) models were trained on ^15^N relaxation data for 400k purely synthetic ‘proteins’ (like other NMR/ML applications) that ranged in size from 40 to 450 ‘amino acids.’ Synthetic dynamics parameters were pseudo-randomly chosen based on typical NMR-derived data. For example, S^2^ values were skewed closer to 1; τ_e_ values were randomly chosen from a distribution between 0 ps and a maximum value that continuously scaled from 1000 ps when S^2^ = 0 to 300 ps when S^2^ = 0.8; and R_ex_ was only allowed for a small percentage of the residues (0 – 20%) and was randomly selected between 1 and 25 s^-1^. These synthetic dynamics parameters were used to generate ^15^N R_1_ and R_2_ relaxation rates and {^1^H}-^15^N heteronuclear NOE values using original model-free formalism plus conformational exchange (i.e., model 4 in the model-free formalism; see Methods). Finally, random errors between 2-8% of the relaxation values were added.

To obtain robust error estimates for the predicted dynamics parameters, a bootstrap ensemble approach was utilized. Ten independent models were trained, each using a different random selection of the training data. Briefly, for each model, the training data was randomized and split into training (90% of the synthetic data) and validation (10%) sets, which were then used for model building. For each residue, the final reported dynamics parameters and their associated errors are calculated as the mean and standard deviation, respectively, of the predictions from the ten models. The ensemble standard deviation represents the epistemic uncertainty (i.e., a measure of the confidence of the DNN based on the noise and/or ambiguity of the training data) of the model. By leveraging the ensemble variance and subsequent calibration (see below), SPINDLE provides fast error estimates and effectively circumvents the need for computationally expensive Monte Carlo simulations traditionally required to estimate fitting errors in data analysis.

### Model Calibration and Synthetic Validation

To validate the DNN models, we generated new ^15^N relaxation datasets for 10k synthetic ‘proteins’ at each static magnetic field using the same approach as described above. We then used SPINDLE to predict the dynamics parameters and compared those to the ground truth values. Initial smaller scale tests suggested that a systematic linear deviation was present for τ_c_ where SPINDLE predicted values lower than the ground truth in a τ_c_ dependent manner. Therefore, we also used the validation datasets to generate linear calibration curves for τ_c_. Local parameters S^2^, τ_e_, and R_ex_ did not require post-hoc calibration.

After calibration, SPINDLE does an excellent job of predicting the global correlation time at all magnetic fields with a mean absolute error (MAE) and root mean square error (RMSE) of ∼0.067 ns and 0.092 ns, respectively (Table 1). This accuracy and precision extend across τ_c_ values ranging from 3 to 35 ns (Figure 2A), as expected from the training data, although the quality of the fits does degrade slightly as the magnetic field increases. The DNN models also accurately predict S^2^, deviating from the ground truth with an overall MAE and RMSE of ∼0.017 and ∼0.027, respectively (Figure 2B and Table 1). For both parameters, Pearson’s correlation coefficients >0.99 are obtained. SPINDLE predicts τ_e_ well, providing values that have MAE and RMSE between 39 – 46 ps and 57 – 65 ps, respectively, from the ground truth (Figure 2C and Table 1). Here, the correlation with magnetic field is the opposite of what was observed for τ_c_ and S^2^ – higher quality fits were obtained for the larger magnetic field strengths (Table 1), similar to what was previously observed in traditional non-linear least squares fitting of NMR relaxation data to model-free parameters by d’Auvergne and Gooley^17^. This phenomenon likely occurs because the high frequency spectral density components of the relaxation rates, especially the R_1_ and heteronuclear NOE, are sampled better at higher static magnetic field strengths, which improves the sensitivity to the τ_e_-dependent term^40,41^. Together, these results demonstrate that SPINDLE can accurately derive fast time scale dynamics information from standard ^15^N relaxation datasets collected at a single magnetic field. Moreover, these errors are well within the typical experimental uncertainties obtained with fitting standard model-free parameters, demonstrating that the accuracy of SPINDLE is similar to traditional least squares fitting methods. Note, all predictions are obtained directly from the relaxation data without the need for initial guesses. Omitting initial guesses eliminates the first step in traditional model-free analysis where an estimate of τ_c_ is calculated from rigid R_2_/R_1_ ratios^42^.

**Table 1.** Validation statistics for SPINDLE DNNs. Each cell gives the mean absolute error (MAE, top value), root mean square error (RMSE, middle value), and Pearson’s correlation coefficient (R_P_, bottom value) for the indicated dynamic parameter and static magnetic field. τ_c_ MAE and RMSD in units of ns. τ_e_ MAE and RMSD in units of ps. R_ex_ MAE and RMSD in units of s-1.

|  | **500**<br>**MHz** | **600**<br>**MHz** | **700**<br>**MHz** | **750**<br>**MHz** | **800**<br>**MHz** | **850**<br>**MHz** | **900**<br>**MHz** | **1000**<br>**MHz** | **1100**<br>**MHz** |
| --- | --- | --- | --- | --- | --- | --- | --- | --- | --- |
| $\tau_c$ | 0.060<br>0.082<br>1.000 | 0.068<br>0.093<br>1.000 | 0.062<br>0.086<br>1.000 | 0.068<br>0.096<br>1.000 | 0.063<br>0.090<br>1.000 | 0.065<br>0.089<br>1.000 | 0.070<br>0.096<br>1.000 | 0.070<br>0.096<br>1.000 | 0.076<br>0.104<br>1.000 |
| $S^2$ | 0.016<br>0.026<br>0.996 | 0.017<br>0.028<br>0.996 | 0.017<br>0.027<br>0.996 | 0.017<br>0.027<br>0.996 | 0.017<br>0.027<br>0.996 | 0.017<br>0.028<br>0.996 | 0.017<br>0.029<br>0.995 | 0.017<br>0.028<br>0.996 | 0.018<br>0.030<br>0.995 |
| $\tau_e$ | 46.123<br>65.541<br>0.942 | 44.761<br>64.179<br>0.945 | 42.193<br>61.786<br>0.949 | 41.344<br>60.921<br>0.950 | 40.575<br>60.049<br>0.952 | 39.922<br>59.289<br>0.953 | 40.184<br>59.128<br>0.953 | 38.475<br>57.607<br>0.956 | 38.687<br>57.418<br>0.956 |
| $R_{ex}$ | 0.284<br>0.759<br>0.992 | 0.333<br>0.873<br>0.988 | 0.352<br>0.929<br>0.987 | 0.358<br>0.951<br>0.986 | 0.362<br>0.971<br>0.986 | 0.398<br>1.044<br>0.982 | 0.433<br>1.120<br>0.980 | 0.457<br>1.171<br>0.979 | 0.511<br>1.330<br>0.973 |

**Figure 2.**
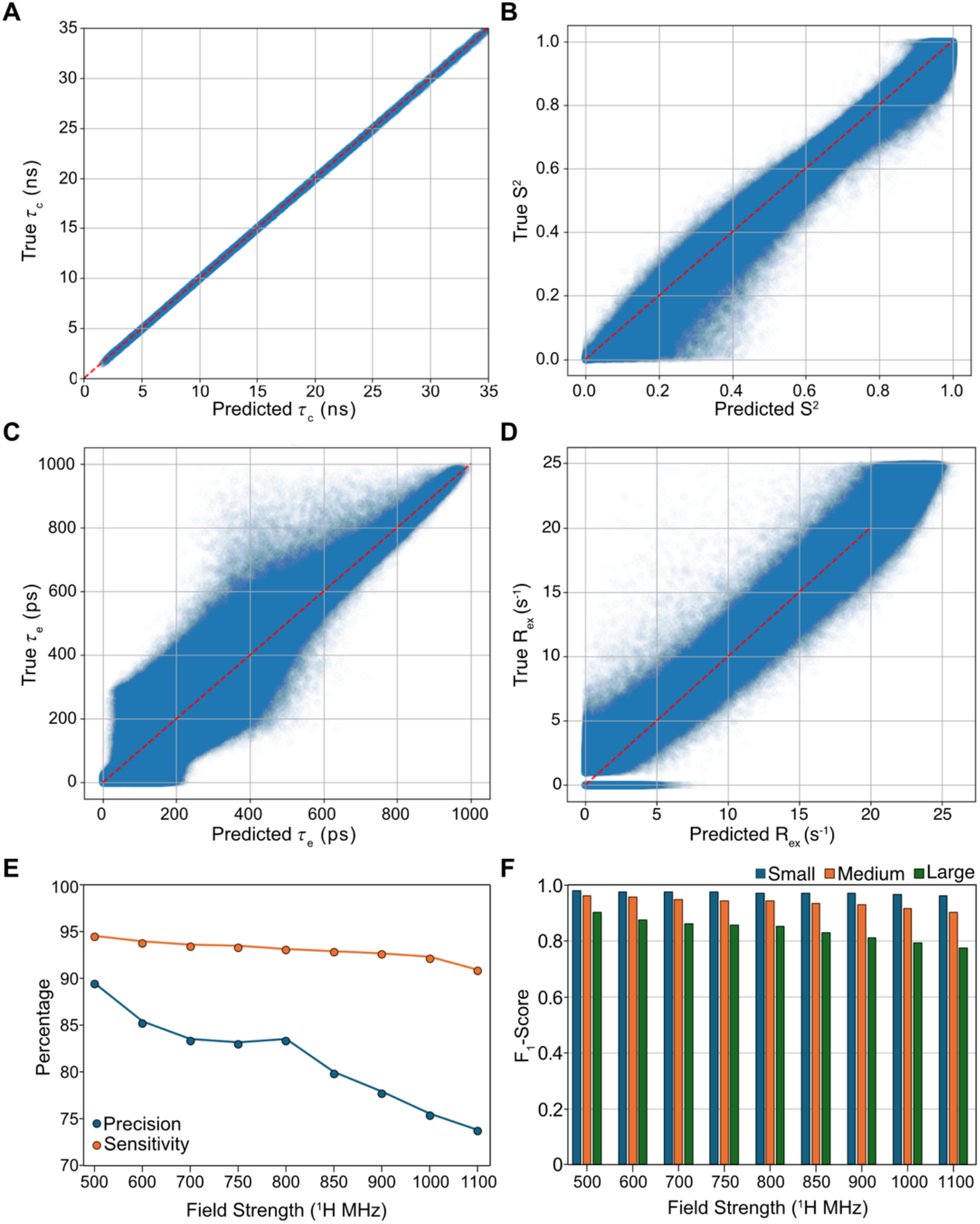
SPINDLE validation on synthetic data. (A-D) Comparison of ground truth (“True”) *vs* SPINDLE-predicted dynamics parameters – (A) global rotational correlation time (τ_c_, in ns), (B) order parameter (S^2^), (C) local effective correlation time (τ_e_, in ps), and (D) chemical exchange contribution (R_ex_, in s^-1^) – for a synthetic validation 500 MHz relaxation dataset for 10,000 ‘proteins’ evaluated using the 500 MHz model. Individual data points are shown in a semi-transparent scatter plot with the red dashed line representing the y=x line. (E) Precision (blue) and sensitivity (orange) of R_ex_ classification as a function of the magnetic field strength. (F) Classification F_1_-scores for R_ex_, grouped by synthetic protein size – small (blue, τ_c_ < 10 ns), medium (orange, 10 ns < τ_c_ < 20 ns), and large (green, τ_c_ > 20 ns) – across the various magnetic field strengths.

Typically, to calculate additional dynamics parameters, such as slow conformational exchange from ^15^N spin relaxation data, additional experimental relaxation datasets are measured at other static magnetic field strengths. In these cases, traditional model selection, based on AIC and/or F-tests using more complicated model-free models, often generates false positives or misses subtle dynamics. We hypothesized that by training the DNN with relaxation data associated with a wide range of possible dynamics parameters SPINDLE would be able to extract additional parameters from the one field data. As seen in Figure 2D and Table 1, SPINDLE can indeed accurately determine the slow conformational exchange R_ex_ parameter from the single field relaxation data. From 500 – 1100 MHz, the MAE and RMSE range between 0.28 – 0.51 s^-1^ and 0.76 – 1.33 s^-1^, respectively, becoming slightly worse at higher magnetic field strengths. We took advantage of the large validation datasets to determine the sensitivity, precision, and false positive rate of SPINDLE for these R_ex_ predictions (Table 2; Figure 2E). That is, how often does SPINDLE predict R_ex_ when it should (i.e., sensitivity), how often does it not predict R_ex_ when it should not (i.e., precision), and how often does it predict R_ex_ when it should not (i.e., false positive rate)? To evaluate more detailed classification metrics, a threshold of R_ex_ ≥ 1.0 s^-1^ was used to define a true exchange event. Regarding sensitivity, SPINDLE finds exchange when it should >91% of the time across all magnetic fields, although the recall is slightly better at low fields than it is at high fields (94.5% at 500 MHz *vs* 90.9% at 1100 MHz; Figure 2E and Table 2). The DNN exhibited a precision of >73.8% (i.e., a false discovery rate of <27%), meaning that when SPINDLE predicted an exchange event, it was correct >73% of the time, again with a strong field dependence (Figure 2E and Table 2). This strong balance between identifying true exchanging residues and minimizing false discoveries resulted in an overall F_1_-score, the harmonic mean of precision and sensitivity (i.e., *F*_1_ = 2 ∗ *Precision* ∗ *Sensitivity*⁄(*Precision* + *Sensitivity*)), of 0.91 – 0.81 (Table 2). Importantly, SPINDLE was highly effective at avoiding false positives, maintaining a low false positive rate between 2.3% - 6.7%; that is, of all the residues that did not have chemical exchange, SPINDLE falsely predicted exchange for only 2.3% to 6.7% of them, again depending on the magnetic field strength. Finally, to further quantitatively evaluate the accuracy of the R_ex_ predictions, we calculated MAE using residues undergoing true chemical exchange with R_ex_ ≥ 1.0 s^-1^; these values ranged between 1.25 s^-1^ – 1.96 s^-1^ for 500 and 1100 MHz field strengths, respectively. Although these MAEs are greater than experimental errors in R_ex_ that can be obtained from relaxation dispersion methods (e.g., CPMG or R_1ρ_), these results demonstrate that SPINDLE can nevertheless identify residues likely undergoing micro-to-millisecond conformational exchange and highlight these for follow up experiments.

**Table 2.**
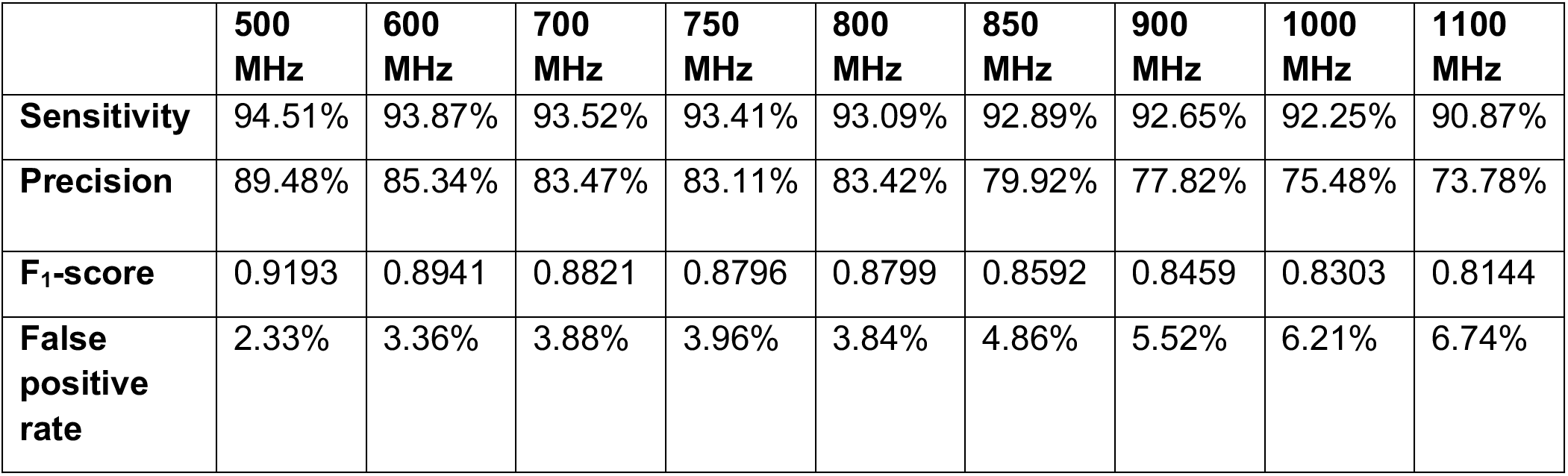
Sensitivity and Precision of SPINDLE for predicting chemical exchange. Each row gives the sensitivity, precision, F_1_-score, or false positive rate for the prediction of chemical exchange (R_ex_) by SPINDLE.

|  | **500**<br>**MHz** | **600**<br>**MHz** | **700**<br>**MHz** | **750**<br>**MHz** | **800**<br>**MHz** | **850**<br>**MHz** | **900**<br>**MHz** | **1000**<br>**MHz** | **1100**<br>**MHz** |
| --- | --- | --- | --- | --- | --- | --- | --- | --- | --- |
| **Sensitivity** | 94.51% | 93.87% | 93.52% | 93.41% | 93.09% | 92.89% | 92.65% | 92.25% | 90.87% |
| **Precision** | 89.48% | 85.34% | 83.47% | 83.11% | 83.42% | 79.92% | 77.82% | 75.48% | 73.78% |
| **F<sub>1</sub>-score** | 0.9193 | 0.8941 | 0.8821 | 0.8796 | 0.8799 | 0.8592 | 0.8459 | 0.8303 | 0.8144 |
| **False**<br>**positive**<br>**rate** | 2.33% | 3.36% | 3.88% | 3.96% | 3.84% | 4.86% | 5.52% | 6.21% | 6.74% |

As noted above, the performance for predicting R_ex_ demonstrated a distinct field dependence, with the overall F_1_-score, for example, decreasing from 0.92 at 500 MHz to 0.81 at 1100 MHz (Table 2). Since ground truth R_ex_ values for each training dataset (based on field strength) for SPINDLE were randomly, uniformly distributed between 1 and 25 s^-1^ (i.e., SPINDLE knows nothing about the field dependence of R_ex_), we hypothesized that the degradation in classification is an artifact of the relative size of the R_ex_ and the intrinsic R_2_. The intrinsic R_2_ is largely governed by both the τ_c_ and S^2^, via dipolar relaxation and the chemical shift anisotropy (CSA), the latter of which scales with the square of the static magnetic field (B_0_^2^). Therefore, larger proteins and higher magnetic fields give rise to larger intrinsic R_2_ rates. For example, a protein with a τ_c_ = 35 ns has a R_2_ = 44.7 s^-1^ and 75.6 s^-1^ at 500 and 1100 MHz, respectively (S^2^ = 1 and τ_e_ = 0 ps). An additional R_ex_ term of 25 s^-1^ represents approximately a third or a quarter of the total R_2_, respectively. The issue is compounded when experimental error is added, here 2%-8% of the total relaxation rate. Thus, the R_ex_ terms, especially at higher magnetic fields, can be overwhelmed by the larger R_2_ and associated noise. To explicitly test this hypothesis, we stratified the classification metrics by protein size using the true τ_c_ (Figure 2F). At 500 MHz, SPINDLE predicts R_ex_ with exceptional accuracy for small proteins (τ_c_ < 10 ns), achieving an F_1_-score of 0.983 (precision of 99.6%); however, for large proteins (τ_c_ > 20 ns), the F_1_-score drops to 0.869 (precision of 81.6%). This size-dependent degradation is exacerbated at 1100 MHz, where the F_1_-score for large proteins drops to 0.730 from 0.947 observed for small proteins (Figure 2F). Importantly, the predictive accuracy for τ_c_ and S^2^ does not degrade with protein size or magnetic field strength (Table 1), since these dynamics parameters are defined by the global interplay of R_1_, R_2_, and NOE. Nevertheless, these results highlight the potential issue of extracting conformational dynamics terms from NMR spin relaxation measurements on large proteins and/or data collected at high magnetic fields.

Finally, to validate that the ensemble standard deviation serves as a proxy for the true statistical error, we used the large validation dataset to evaluate the statistical calibration of the DNN ensemble. First, we asked how often the ground truth value falls within the 95% confidence interval of the SPINDLE predicted value. For the validation data generated at 600, 800, and 1100 MHz, we found that at all three fields ∼97% (τ_c_), ∼61% (S^2^), ∼47% (τ_e_), and ∼32% (R_ex_) of the time the true value was within the 95% confidence interval. As is common in deep learning ensembles, the absolute magnitude of the epistemic uncertainty (i.e., uncertainty in the models) tends to be under calibrated compared to the true residual error, as the models frequently converge to highly similar predictions. To further evaluate the epistemic uncertainty of the ensemble, we calculated the Pearson’s correlation coefficient (R_P_) between the standard deviation of the ensemble predictions and the true residual error. The positive correlations (e.g., R_P_ ∼0.4 for τ_c_ and S^2^, R_P_ ∼0.2 for τ_e_, and R_P_ ∼0.6 for R_ex_) demonstrated that ensemble variance acts as a relative indicator of prediction error. To provide robust statistical bounds for standard structural/biophysical analysis, the raw ensemble standard deviations were scaled by parameter-specific multipliers determined on the validation set, thereby ensuring that the 95% confidence intervals accurately achieve 95% coverage probability across the entire parameter space. Multipliers of ∼3.9–3.1 for S^2^, and ∼6.7–5.7 for τ_e_ and ∼3.4–2.9 for R_ex_ were derived to inflate the raw standard deviations to a true 95% coverage probability. Conversely, the ensemble was slightly underconfident regarding its predictions of τ_c_ (requiring multipliers of ∼0.8–1.0). By applying these parameter-specific calibration multipliers, SPINDLE provides end-users with absolute, statistically rigorous 95% confidence intervals at a fraction of the computational cost of traditional Monte Carlo simulations.

### SPINDLE Outperforms Existing model-free Analysis Software

Several programs or wrappers exist for extracting relaxation parameters using model-free formalism introduced by Lipari and Szabo^16,18–20^. These programs allow users to fit their data to one of the five (or more) models used in model-free using non-linear least squares-based optimizers. Here, we tested the efficacy of SPINDLE against one of these programs, *modelfree v4* (ref ^16^), using four *in silico* ^15^N relaxation datasets of 1000 ‘proteins’ generated with either model 2 (global τ_c_ and local S^2^ and τ_e_) or model 4 (global τ_c_ and local S^2^, τ_e_, and R_ex_) at 800 MHz and 600 MHz. The relaxation parameters for these synthetic relaxation datasets were extracted with either the corresponding model using *modelfree* or SPINDLE and then compared to the ground truth values.

Overall, SPINDLE outperformed *modelfree* in both computational efficiency and predictive accuracy and precision. For each of the four datasets, SPINDLE needed ∼50 minutes on a CPU to predict the dynamics parameters for the 1000 ‘proteins.’ On the other hand, *modelfree* required exponentially longer computational time: model 2 datasets needed ∼7500 minutes, whereas model 4 datasets needed ∼21,000 minutes. Note, *modelfree* was executed using the GNU *parallel* command to minimize total run time/maximize CPU usage on a NMRbox server, meaning that the actual run times were shorter than the cumulative computation times reported here. Regardless, SPINDLE consistently completed analyses much faster than *modelfree* for both the cumulative computation time and actual time.

Additionally, *modelfree* failed to converge for many proteins. Of the 1000 proteins in the model 2 datasets, ∼450 runs were completed successfully for datasets at each magnetic field. This failure rate was a combination of a size limitation within the *modelfree* software (<300 residues; 60% of the failed runs) and a general failure of the calculation to converge (40% of the failed runs)^17,40,43^. Worse, in the model 4 datasets, *modelfree* successfully calculated dynamics parameters for only 40 and 26 datasets at 800 MHz and 600 MHz, respectively, before the process timed out (i.e., the calculation took >45 minutes). For the model 4 datasets, this failure of convergence was expected as too little experimental data (i.e., only single-field relaxation data for four fitted parameters) was given to *modelfree* to determine dynamics parameters by non-linear least squares fitting^17,40,43^.

We then compared the accuracy of the extracted dynamics parameters for the model 2 and model 4 datasets from the two analysis pipelines. Although the relaxation parameters determined from the model 2 datasets using *modelfree* were good, SPINDLE demonstrated better accuracy and precision to the ground truth values at both fields when comparing the RMSE values (Figure 3 and Supplemental Figure 1). Not surprisingly, *modelfree* performed much worse at fitting the more complicated model 4 datasets for both fields than the simpler model 2 datasets (Figure 4 and Supplemental Figure 2). For the calculations that converged, a big issue for *modelfree* involved the determination of R_ex_; here, the RMSE values (3.97 and 10.93 s^-1^ for 600 and 800 MHz, respectively) approached or exceeded the maximum value of R_ex_ (10 s^-1^) used to generate the data. At 800 MHz, because of the issues with R_ex_, *modelfree* generally underfit τ_c_ and τ_e_ and poorly fit S^2^ (Supplemental Figure 2B). The performance of *modelfree* ‘improved’ at 600 MHz, where the S^2^ and τ_e_ values were generally more accurate, but R_ex_ remained overfit and τ_c_ underfit (Figure 4B). Although the low RMSEs for S^2^ and τ_e_ for the 600 MHz dataset imply accurate fitting, they mask the fact that only 4% of the ‘proteins’ in the dataset were fit. In comparison, SPINDLE had consistent results across all the parameters with strong fits for all 1000 ‘proteins’ at both fields for S^2^, R_ex_, and τ_c_ (Figure 4A and Supplemental Figure 2A).

**Figure 3.**
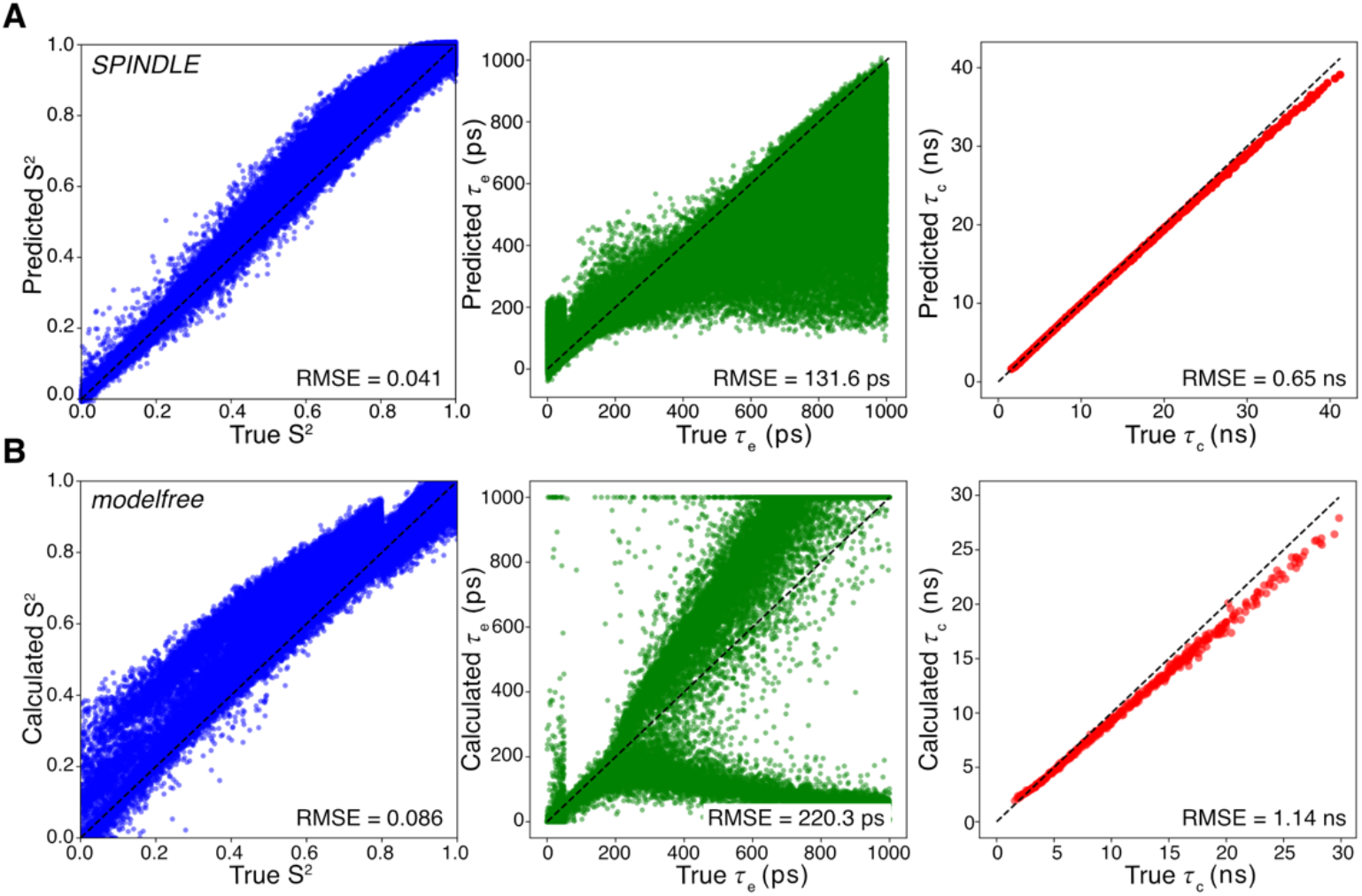
Comparison of SPINDLE predictions and conventional non-linear least squares analysis on synthetic 600 MHz model 2 dynamics data. Comparison between (A) SPINDLE deep learning predictions and (B) traditional non-linear least-squares optimization (via *modelfree* v4) using synthetic model 2 (i.e., global τ_c_ and local S^2^ and τ_e_) relaxation data generated for 1000 ‘proteins’ at 600 MHz. *Left column*, predicted/calculated *vs* ground truth S^2^; *middle column*, predicted/calculated *vs* ground truth τ_e_; and *right column*, predicted/calculated *vs* ground truth τ_c_. In all plots, the black dashed line indicates the y=x line, and the root mean square error (RMSE) value comparing the predicted/calculated and ground truth values are reported in the lower-right corner of each plot.

**Figure 4.**
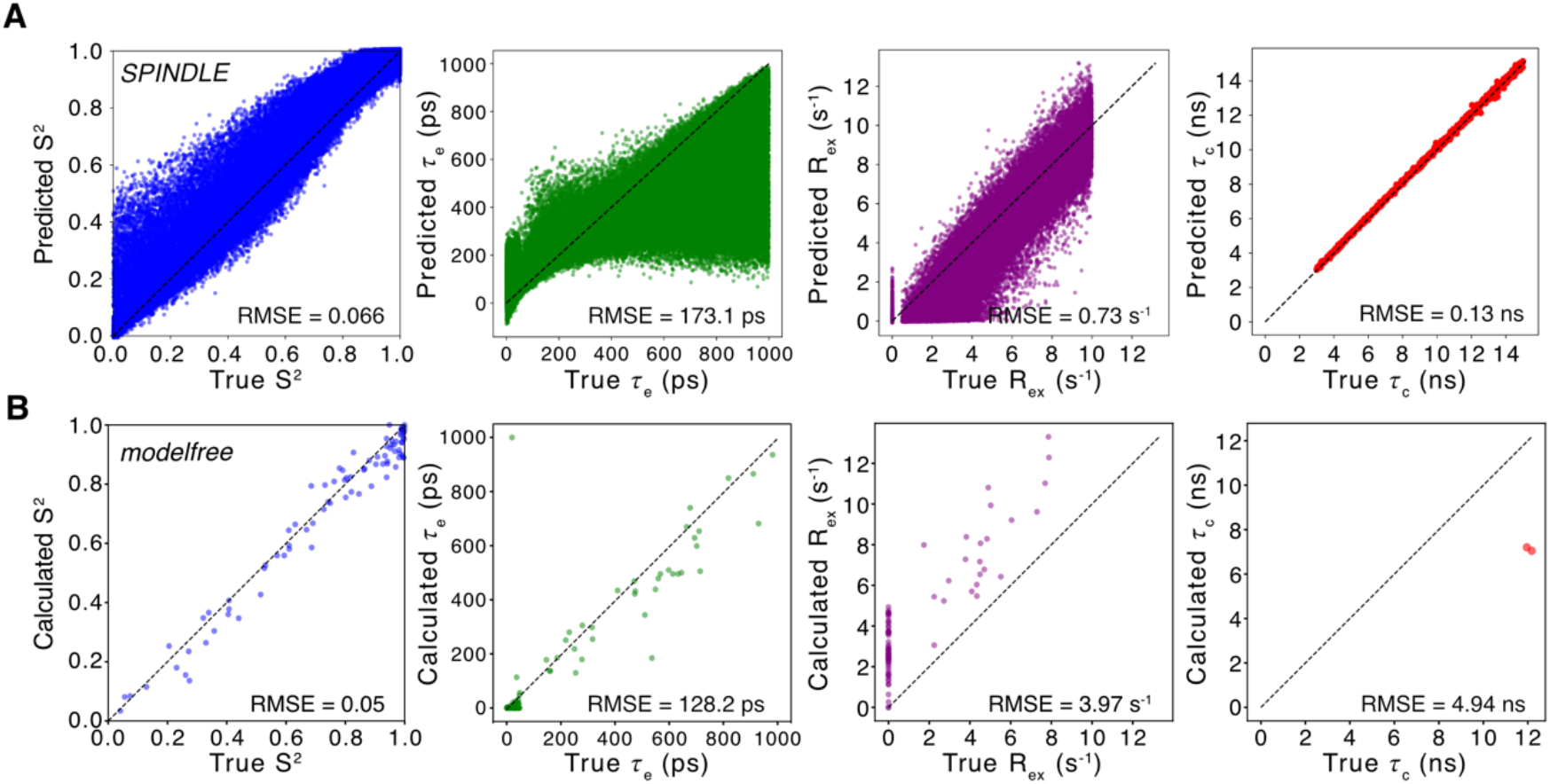
Comparison of SPINDLE predictions and conventional non-linear least squares analysis on synthetic 600 MHz model 4 dynamics data. Comparison between (A) SPINDLE deep learning predictions and (B) traditional non-linear least-squares optimization (via *modelfree* v4) using synthetic model 4 (i.e., global τ_c_ and local S^2^, τ_e_, and R_ex_) relaxation data generated for 1000 ‘proteins’ at 600 MHz. *First column*, predicted/calculated *vs* ground truth S^2^; *second column*, predicted/calculated *vs* ground truth τ_e_; *third column*, predicted/calculated *vs* ground truth R_ex_; and *forth column*, predicted/calculated *vs* ground truth τ_c_. In all plots, the black dashed line indicates the y=x line, and the root mean square error (RMSE) value comparing the predicted/calculated and ground truth values are reported in the lower-right corner of each plot.

The above comparisons do not mimic the actual process of model-free data analysis. That is, a user would not typically fit every residue to a single model but would systematically test each residue for the various models and select the appropriate model based on hypothesis testing^16^ or AIC^17^. To mimic a more standard workflow, we used the *modelfree* wrapper *FAST-Modelfree*^20^ to individually fit the 600 and 800 MHz *in silico* model 4 relaxation datasets. *FAST-Modelfree* automates the process of fitting each residue to all five model-free models and selects the optimal model using F-tests and sum-of-squared-error criteria^20^. Like *modelfree*, *FAST-Modelfree* was executed using GNU *parallel*. *FAST-Modelfree* required ∼220,000 minutes to run the model 4 800 MHz and 600 MHz datasets. This represents an enormous increase in computational cost compared to SPINDLE, which required only ∼50 minutes for the same datasets. Even with an increased timeout time (600 minutes), *FAST-Modelfree* still failed to fit every residue, excluding ∼30% of the residues in each dataset. This outcome was not completely unexpected as *FAST-Modelfree* has previously been reported to exclude residues during fitting^20^. Excluded residues could have resulted from setting fitting thresholds too high, adding too much error, or the program not converging. In contrast, SPINDLE predicts relaxation parameters using a fundamentally different approach and therefore does not exclude residues in the manner observed with *FAST-Modelfree*.

Like *modelfree*, dynamics parameters calculated by *FAST-Modelfree* had much lower accuracy compared to SPINDLE and the graphs show extremely weak precision and accuracy to the ground truth values (compare Figure 4A to 5A and Supplemental Figure 2A to 3A). Most worrisome, inspection of the assigned models revealed that *FAST-Modelfree* incorrectly classified a substantial portion of residues as model 5 (global τ_c_ and local S_f_^2^, S_s_^2^, and τ_s_) even though the datasets contained no model 5 data (Figure 5B and Supplemental Figure 3B). In fact, a large portion of the incorrectly assigned model 5 residues originated from data generated to represent model 2, which contains only S^2^ and τ_e_ values. In conclusion, we show here that SPINDLE is a highly accurate, precise, and fast alternative to standard programs used for model-free analysis.

**Figure 5.**
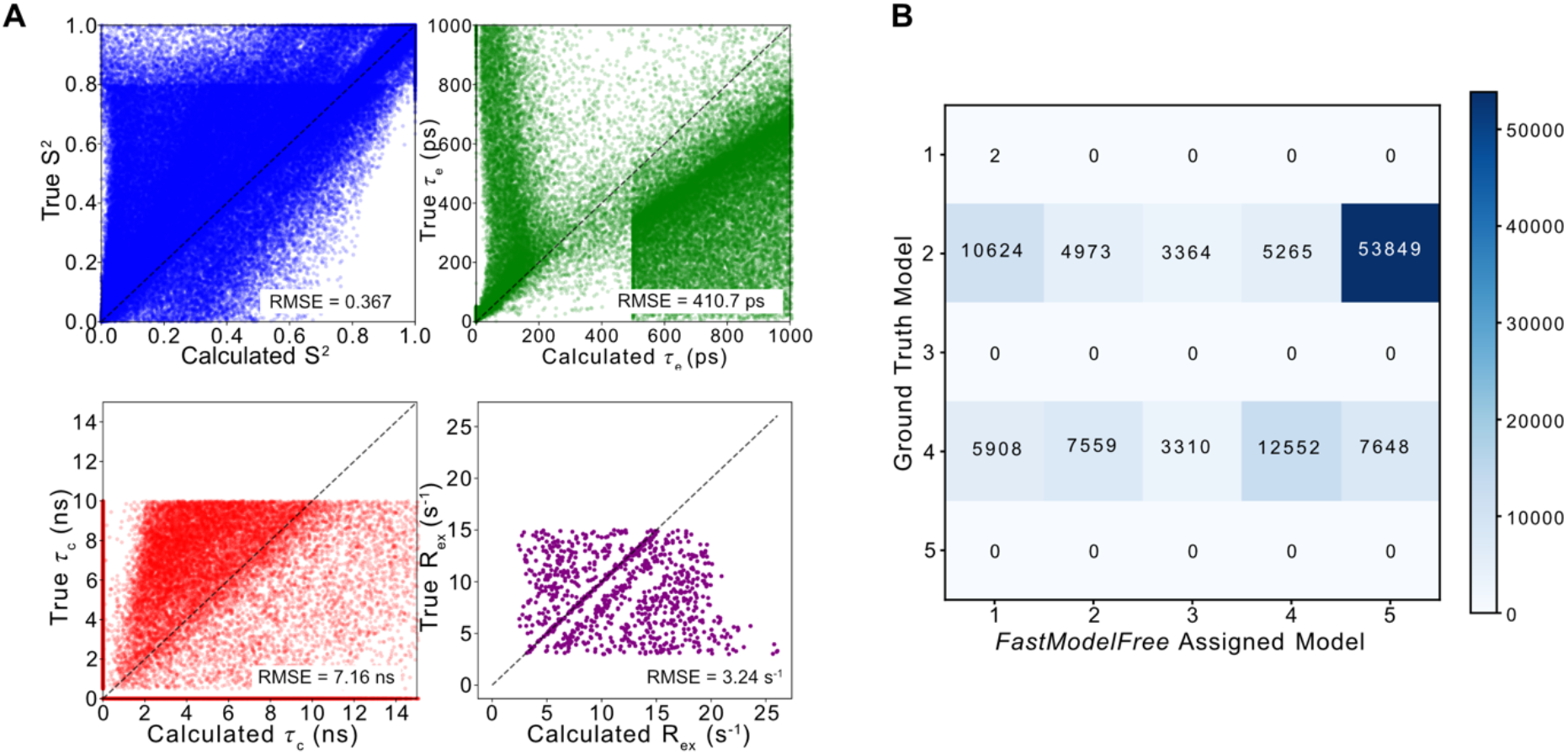
Evaluation of automated model selection on synthetic 600 MHz relaxation data. (A) Comparison of ground truth *vs* non-linear least-squares calculated (via *FAST-modelfree and modelfree* v4) dynamics parameters from relaxation data generated for 1000 synthetic proteins at 600 MHz. *Top-left*, calculated *vs* ground truth S^2^; *top-right*, calculated *vs* ground truth τ_e_; *bottom-left*, calculated *vs* ground truth τ_c_; and *bottom-right*, calculated *vs* ground truth R_ex_. In all plots, the black dashed line indicates the y=x line, and the root mean square error (RMSE) value comparing the calculated and ground truth values are reported in the lower-right corner of each plot. (B) Heatmap comparing the actual model-free model selection of the synthetic ground truth data (y-axis) against the model selected by hypothesis testing within *FAST-modelfree* (x-axis). The numbers within each cell indicate the total number of assigned residues with the color scale corresponding to the residue count.

### Validation with Experimental Data

To further test SPINDLE, the analysis pipeline was evaluated using experimental ^15^N relaxation datasets from the BMRB. Selected datasets contained amide ^15^N R_1_, R_2_, and heteronuclear NOE relaxation data, along with the previously determined dynamics parameters. 51 of the 76 relaxation datasets in the BMRB, which contained 71 different relaxation datasets from different field strengths and/or conditions, were ultimately included in the analysis. Since the global tumbling time is not a reported value in the BMRB, τ_c_ values were retrieved from the corresponding publications.

SPINDLE predicted τ_c_ values for the experimental relaxation data ranging from 3.8 ns to 38 ns with high precision (Figure 6A; R_P_ = 0.947). The precision over this range was slightly unexpected as the model was trained on synthetic datasets with τ_c_s between 5 ns and 35 ns, yet the DNN faithfully extrapolated outside of this range. Surprisingly, SPINDLE does an excellent job of capturing the isotropic τ_c_s reported for axially symmetric and anisotropic models (red points in Figure 6A) even though an isotropic tumbling model was used to generate the training data. The models predicted the tumbling times of small peptides such as the 34-residue long c-di-GMP-sequestering peptide (BMRB ID: 50001) and the 42-residue long WW3 domain of hNedd4-1 (BMRB ID: 18971) to be 3.57 ns and 4.19 ns, respectively, whereas the reported τ_c_ values were 3.8 ns and 3.9 ns, respectively. The τ_c_s of large complexes were also accurately predicted by SPINDLE. As an example, the global tumbling time of the 110 kDa hexamer inorganic pyrophosphatase was predicted by SPINDLE to be 39.6 ns compared to the reported value of 37.9 ± 0.3 ns (BMRB ID: 50212). Correctly fitting the global tumbling time for this hexamer demonstrates not only the broad range of τ_c_s SPINDLE can predict, but also that the model was not directly correlating protein length to τ_c_. Instead, SPINDLE recognized the hexamer as a large, slowly tumbling complex rather than fitting the τ_c_ for one of its six 18.3 kDa monomers. Additional evidence that SPINDLE does not rely solely on the number of residues when predicting τ_c_ was observed with the 22.5 kDa human growth hormone (rhGH; BMRB ID: 4689). SPINDLE predicted τ_c_s of 12.74 ns and 14.41 ns for pH 2.7 and pH 7 conditions, respectively, which compare favorably to the reported values of 11.9 ns and 13.9 ns. Both τ_c_s are slower than what would be expected for a protein of rhGH’s size using a spherical model (theoretical τ_c_ of 9 ns). The authors hypothesized that the slower global tumbling could be due to the propensity of rhGH to aggregate^44^. Similarly, SPINDLE accurately predicted τ_c_s for wildtype and mutant forms of a neuropeptide bound to dodecylphosphocholine (DPC) micelles to be 11 ns and 10.27 ns, respectively, whereas the reported values were 8.96 ± 0.10 ns and 8.22 ± 0.09 ns (BMRB IDs: 5550 and 5549). Finally, to illustrate that SPINDLE does not just work in the extremes, the DNN models were able to accurately predict τ_c_s for human ubiquitin (8.6 kDa) of 4.67 ns and 4.50 ns for 600 MHz and 500 MHz relaxation data, respectively, which compare extremely well with the reported value of 4.6 ns obtained from simultaneously fitting the two field relaxation data (BMRB ID: 4245). Together, these results demonstrate that SPINDLE does not correlate global tumbling time with the number of residues; that even though SPINDLE was trained on an isotropic model for global tumbling, it captures τ_c_ from data previously fit with isotropic and non-isotropic models; and that SPINDLE can accurately predict τ_c_s from relaxation data collected for a given protein under differing experimental conditions (e.g., pH, temperature, mutation, or bound state).

**Figure 6.**
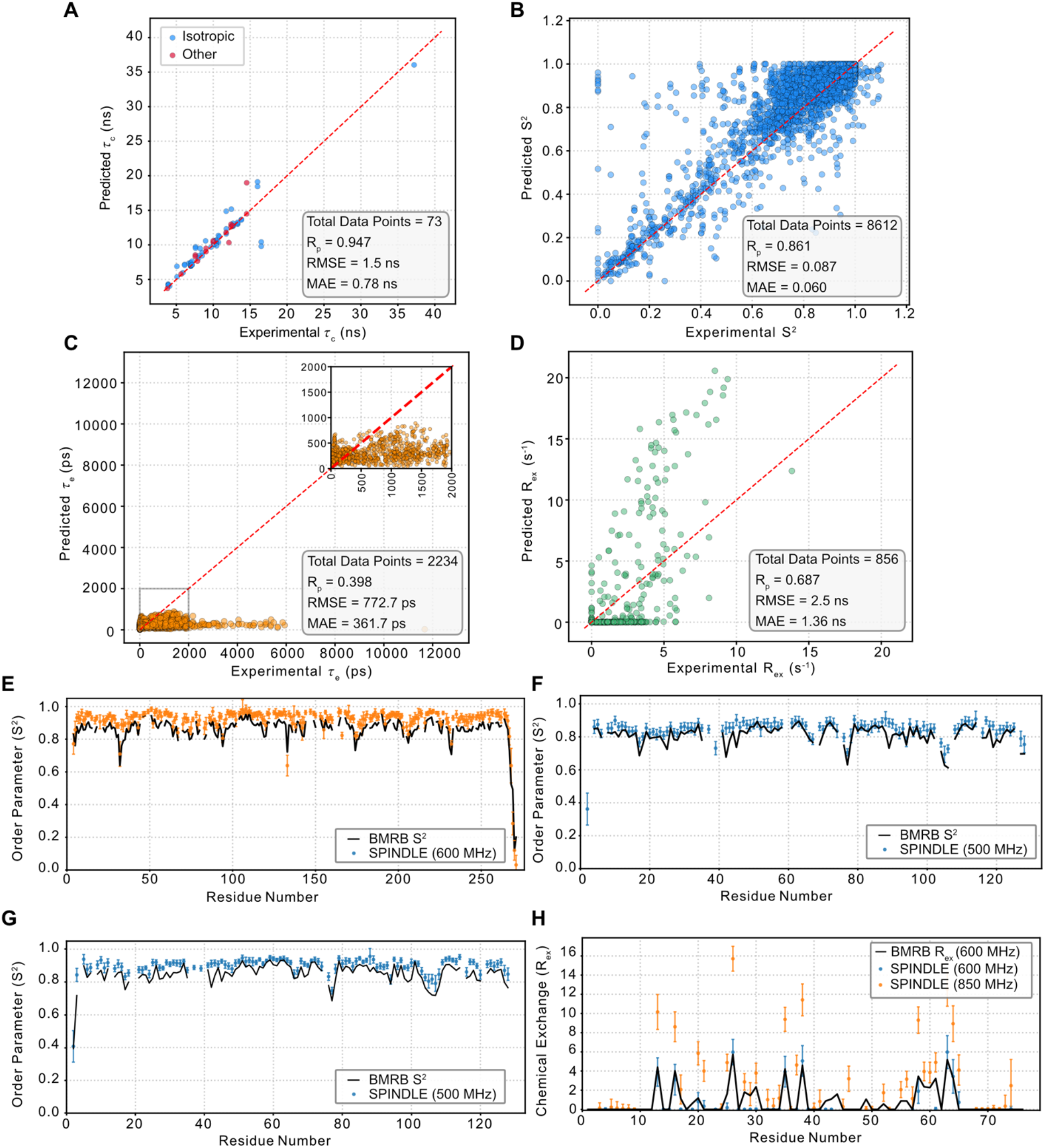
Performance of SPINDLE on experimental NMR relaxation data. (A-D) Correlation between SPINDLE predicted and published model-free parameters deposited in the BMRB. The global τ_c_, color-coded by the reported global diffusion tensor, is shown in (A); the order parameter is shown in (B); the internal correlation time τ_e_ is shown in (C) with an inset displaying an expanded view of 0 – 2000 ps; and the R_ex_ exchange contribution is shown in (D). For each panel, the number of compared proteins/residues is provided along with a summary of the agreement between the two methods. The red dashed line denotes the y=x. (E-H) Per-residue comparisons of published model-free parameters (solid black lines) and SPINDLE predictions (colored markers with error bars). (E) shows the comparison of S^2^ values derived from 600 MHz relaxation data reported in BMRB ID 6838. (F) and (G) show the comparison for S^2^ values derived for 500 MHz relaxation data collected at 16 °C and 44 °C as reported in BMRB ID 6243. (H) shows the comparison of R_ex_ exchange contribution at 600 MHz compared to the SPINDLE predictions from the 600 MHz (blue points) and 800 MHz (orange points) relaxation data reported in BMRB ID 27011.

SPINDLE also demonstrated excellent performance in predicting order parameters (Figure 6B). In general, there was a strong correlation (R_P_ = 0.861) between the SPINDLE-predicted and BMRB-reported values across the full range of S^2^. Using single field data, the DNN models were able to precisely predict order parameters from data with multiple fields, as exhibited with the S^2^ values for the 29.5 kDa β-lactamase PSE-4 collected at 500, 600, and 800 MHz (BMRB ID: 6838; Figure 6E and Supplemental Figure 4A). As with τ_c_, SPINDLE precisely predicted order parameters for multiple conditions including temperature and pH, as demonstrated with its predictions of S^2^ for the 14 kDa azurin at 16 °C and 44 °C (BMRB ID: 6243; Figure 6F and 6G and Supplemental Figure 4B and 4C), and the S^2^ for the 22.5 kDa rhGH, at pH 2.7 and 7 (BMRB ID: 4689; Supplemental Figure 5A and 5B). In addition, SPINDLE accurately predicted variations in S^2^ values for the wildtype and mutant forms of a neuropeptide bound to DPC micelles (BMRB IDs: 5549 and 5550; Supplemental Figures 5C and 5D), as well as galectin-3C bound to three different carbohydrate substrates (BMRB IDs: 50283, 50284, and 50285; Supplemental Figure 6), further supporting the conclusion that SPINDLE can reliably predict changes in order parameters due to environmental or conformational changes. Finally, protein size continued to have minimal impact on DNN performance, as SPINDLE S^2^ predictions for both the 110 kDa hexamer inorganic pyrophosphatase (BMRB ID: 50212; Supplemental 7A) and the 34-residue long c-di-GMP-sequestering peptide (BMRB ID: 50001; Supplemental 7B) agreed very well with the previously reported values.

In contrast to τ_c_ and S^2^, SPINDLE tended to underfit the timescale of fast internal motion as seen in the comparison of the predicted and reported τ_e_ values (Figure 6C). A large fraction of the τ_e_ values were not being captured by SPINDLE resulting in a poor correlation when comparing all the data together (R_P_ = 0.398). One source of this underfitting could be because the model was only trained with τ_e_ values up to 1000 ps, ensuring that ωτ_e_ << 1 (where ω is the frequence in rad s^-1^), and many of the BRMB-derived values go far beyond this value. In fact, as seen in Figure 6C, several studies report τ_e_ values >5000 ps, which would no longer be in the fast-tumbling limit as defined by Lipari and Szabo^8,9^, and in some cases, these reported τ_e_ values started approaching the global tumbling time. Furthermore, the discrepancy in Figure 6C could result from known computational issues of traditional non-linear least squares fitting methods for model-free parameters. First, the non-linear optimization landscape of the parameter space contains long, flat valleys where local optimization algorithms suffer from gradient loss which causes τ_e_ to artificially diverge towards infinity or merge with τ_c_ (ref ^40,43^). As demonstrated by d’Auvergne and Gooley, when model elimination protocols are omitted prior to model selection, these failed optimizations frequently yield lower minimization errors (i.e., χ^2^) than the true model, leading in unrealistic τ_e_ values (>1000-5000 ps)^43^. Additionally, as described above, NMR relaxation data becomes less sensitive to τ_e_ near the extreme narrowing limit (i.e., ω^2^τ^2^ << 1) or for highly rigid residues (i.e., S^2^ ≍ 1) because the contribution of τ_e_ to the spectral density function can be hidden by the experimental noise. Thus, the calculated τ_e_ values under these conditions can have propagated uncertainties that span several hundred picoseconds^41,45^.

As we observed with the synthetic datasets, SPINDLE was able to reasonably predict µs-ms timescale chemical exchange values from experimental single field relaxation data (Figure 6D). A moderate correlation (R_P_ = 0.687) was seen between all the reported R_ex_ values and those predicted by SPINDLE. SPINDLE also correctly identified the field dependence of R_ex_, as seen with the higher R_ex_ rates at 850 MHz compared to those at 600 MHz for the domain factor 1.1, a part of an RNA polymerase (BMRB ID: 27011; Figure 6H). However, the differences between the two chemical exchange rates slightly exceeded the expected factor of two (i.e., the square of the ratio of the two fields), suggesting that SPINDLE tends to slightly overfit chemical exchange contributions at high fields as described above. Despite this, the chemical exchange at each field had little effect in changing how accurate the global tumbling times were for both fields, as the SPINDLE τ_c_s were 7 ns and 7.4 ns for 600 MHz and 850 MHz respectively, while the reported value was 6.13 ns. This example further demonstrates the ability of SPINDLE to identify R_ex_ terms from single field relaxation data, and based on these results, we would suggest follow-up experiments with more accurate and informative relaxation dispersion methods.

As an aside, the use of the BMRB for relaxation data analysis was plagued by a variety of problems with entries in the database. For example, situations were encountered where the incorrect units were given for the relaxation rates (e.g., data were in s or ms^-1^ instead of the indicated s^-1^), the incorrect field strength or sample condition was listed for one or more of the entries, or simply the incorrect data was deposited. If the BMRB is to be used as a possible resource for future machine learning endeavors or more simply as a resource for FAIR – Findable, Accessible, Interoperable, and Resuable – digital data management, better care has to be taken by the community and BMRB to ensure accurate and precise deposition of the data.

### SPINDLE Explainability and Ambiguity

To understand the internal decision-making process of SPINDLE and transition the algorithm from a ‘black box’ to a potential interpretable structural biology tool, we took advantage of the architecture of the model to implement a DNN explainability feature. This feature relies on a retroactive extraction of the self-attention layer to map which inputs (i.e., sequence specific ^15^N relaxation data) the DNN determined were important for τ_c_ prediction. By storing the internal attention weight matrices generated within the transformer block during prediction, we can directly quantify the relative influence that the relaxation profiles for each residue has on the final prediction. Thus, this explainability feature gets to the core of how SPINDLE works, which is predicting four dynamics parameters from three experimental observables. We hypothesized that the self-attention layer would down weight dynamic residues and emphasize rigid residues, because those relaxation rates would reflect global τ_c_ more than local internal dynamics.

To test this hypothesis, we initially generated synthetic relaxation datasets at 600 and 800 MHz and predicted the dynamics parameters with SPINDLE. When the residue specific attention/importance scores are plotted with S^2^ or R_ex_, we observed larger attention for residues with larger S^2^ and smaller R_ex_ values, supporting our hypothesis (Supplemental Figure 8). We next extracted the residue specific attention scores for experimental data from the BMRB (Figure 7). For most of the experimental datasets, the SPINDLE attention scores mimic what was observed for the synthetic data (Figure 7A-C), where more importance for predicting τ_c_ was placed on residues with high S^2^ and no R_ex_. Thus, in these cases, the self-attention layer within the DNN acts very much like a human would in this analysis by filtering out flexible and focusing on rigid residues, as hypothesized. We noted, however, several cases, especially when the model predicts many residues with R_ex_, where high attention scores are given to residues at the termini with very low S^2^ values (e.g., Figure 7D). This case makes it clear that the attention layer within the DNN prioritizes the absence of µs-ms chemical exchange over flexibility on the ps-ns timescale. In these cases, the attention layer is utilizing a different strategy: either identifying outliers to penalize in the subsequent global pooling layer or using these residues to get good values of τ_e_ which can then be removed from the reduced correlation time (τ) in the second term of the spectral density function (J(ω)) to give good predictions for τ_c_. These results demonstrate that in many cases SPINDLE has learned to focus on rigid residues much like a human analyst while also learning that in certain instances flexible residues still contain valuable information for determining τ_c_.

**Figure 7.**
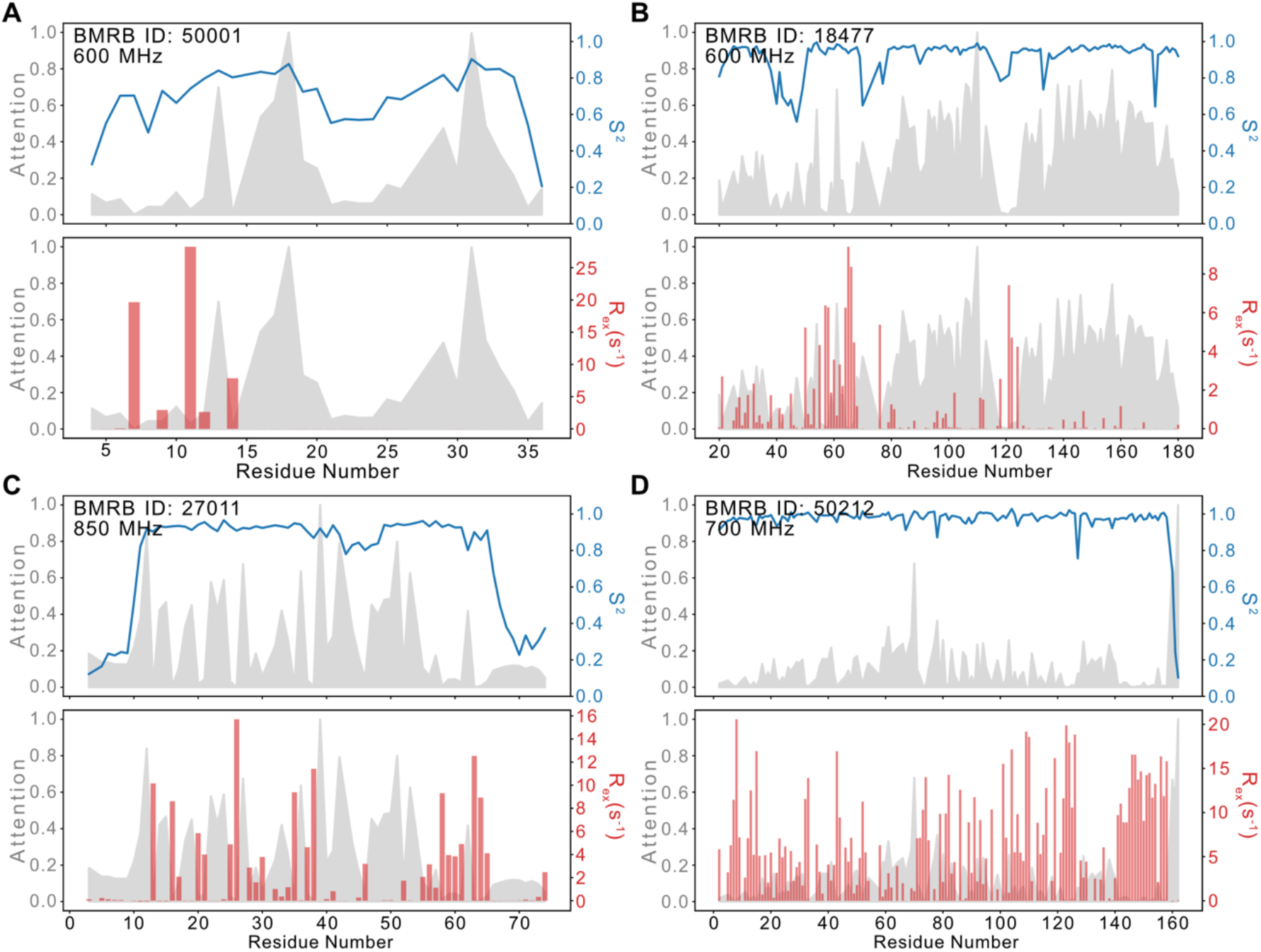
SPINDLE self-attention profiles highlight which residues are important for extracting dynamics parameters. (A-D) Normalized per-residue attention weights extracted from the SPINDLE Multi-Head Self-Attention layer (grey shaded area, left y-axis) overlaid with predicted S^2^ (blue lines, top subplot, right y-axis) and R_ex_ (red bars, bottom subplot, right y-axis). Attention weights and predicted dynamics parameters were obtained from experimental ^15^N relaxation data for BMRB IDs (A) 50001, (B) 18477, (C) 27011, and (D) 50212.

Beyond providing a glimpse into the rationale for the predictions made by the model, the self-attention mechanism offers a novel framework for identifying allosteric networks from NMR relaxation data. To realize this application, the architecture of SPINDLE needs to be adapted to accept multiple relaxation datasets simultaneously (e.g., apo *vs* ligand bound or wildtype *vs* mutant constructs) and retrained on the multi-state dynamic profiles. Because allosteric communication can alter local ps-ns motions (S^2^) or µs-ms conformation conformational exchange, cross-attention and self-attention layers could possibly track how perturbations alter the dynamic landscape across a protein’s structure. Therefore, with this expanded multi-input structure and self-attention mechanism, this DNN strategy could provide an automated, unbiased method to identify allosterically coupled residues and visualize how dynamic perturbations propagate through a protein structure. We are actively pursuing this line of research in ongoing work.

Unlike traditional model-free analysis pipelines that utilize discrete model selection protocols (e.g., AIC or hypothesis testing) to eliminate unresolved dynamics parameters and help with degeneracy, SPINDLE maintains the full parameter space and reports these mathematical degeneracies via the calibrated ensemble variance. We therefore define an “ambiguity” threshold to describe the mathematical degeneracy where single-field experimental ^15^N spin relaxation data are insufficient to provide constraints needed to resolve all four dynamic variables. To calibrate ambiguity thresholds for S^2^, τ_e_, and R_ex_, we first defined error ceilings of 0.10, 150 ps, and 2.5 s^-1^ for S^2^, τ_e_, and R_ex_, respectively. These ceilings define the maximum absolute error that can be tolerated before the prediction becomes misleading. We then went back to the 10k synthetic ‘protein’ validation test datasets, calculated the true absolute error for every prediction, and determined if true absolute error was greater (i.e., flagged ambiguous) or less (i.e., good fit) than these ceilings. Finally, we used this classification and the calibrated standard deviation of the DNN models to generate precision-recall curves to calibrate the ambiguity threshold. This procedure was conducted for each field, resulting in similar ambiguity thresholds for each parameter (∼0.03 for S^2^, ∼24 ps for τ_e_, and ∼0.5 s^-1^ for R_ex_). Therefore, in the SPINDLE pipeline, if a dynamics parameter has a corrected error greater than the average ambiguity threshold, the fit is flagged indicating that the predicted value may be unreliably predicted by the DNN. Ultimately, this parameter-specific ambiguity labeling prevents the over-interpretation of the relaxation data, while preserving high-confidence predictions. For the end user, the ambiguity flag replaces false certainty of automated model selection with transparency by explicitly showing the information boundaries of the single-field data.

## Conclusion

Herein, we describe SPINDLE, a new DNN tool for predicting model-free dynamics parameters, which describe fast (ps-ns; τ_c_, S^2^, and τ_e_) and slow (µs-ms; R_ex_) timescale motions, from NMR ^15^N relaxation data collected at a single static magnetic field. By training an ensemble of DNN models on a large dataset of synthetic ‘proteins’ encompassing a wide dynamics space, we enabled SPINDLE to accurately predict these model-free dynamics parameters and their associated errors from single-field NMR spin relaxation data in a fraction of the time of existing analysis software with minimal intervention by the user regarding dynamical model selection. Further, we extensively used independent testing datasets to not only calibrate the predictions of the models but also determine their standard deviation and then define an ambiguity threshold to provide additional confidence in the DNN predictions. We also used synthetic datasets to show that SPINDLE out-performs existing non-linear least squares-based analysis pipelines and that the hypothesis-based selection criteria that is widely used for model-free model selection is prone to bad model selection. Lastly, SPINDLE recapitulates the dynamics parameters calculated from ∼50 experimental datasets contained in the BMRB, predicting similar τ_c_ and S^2^ values across a wide range of protein size, condition, and tumbling models (i.e., isotropic and axially symmetric) and in many cases similar R_ex_ values. Finally, we take advantage of the internal workings of the DNN (i.e., the multi-head self-attention layer) to extract attention scores, which report on the relative importance of a residue’s relaxation data in the prediction of dynamics parameters. Overall, we anticipate that SPINDLE will expand the use of NMR spin relaxation data and model-free analysis by making this pipeline easier, faster, and available to labs with limited magnet time and/or sample stability.

## MATERIALS AND METHODS

### Training data generation

Training deep learning networks requires a large set of features (i.e., inputs – ^15^N relaxation rates) and their corresponding labels (i.e., outputs – τ_c_, S^2^, τ_e_, and R_ex_). SPINDLE was trained exclusively on synthetic data as described below. At each magnetic field, ^15^N relaxation rates were generated for 400k synthetic ‘proteins’ ranging from 40 to 451 residues. To ensure equal coverage across this size range, proteins were stratified into small (40 – 140 ‘amino acids’), medium (141 – 240 ‘amino acids’), large (241 – 340 ‘amino acids’), and very large (341 – 451 ‘amino acids’) protein bins, with 100k proteins generated for each bin. Within a size bin, a random number of amino acids was chosen, and a base rotational correlation time (τ_c_) was calculated as *τ_c_*_,*base*_ = 0.0005998 ∗ *n_residues_* ∗ 110 + 0.1674, where 0.0005998 and 0.1674 are the slope and intercept to a linear correlation between molecular weight and τ_c_, respectively (https://nesgwiki.chem.buffalo.edu/index.php/NMR_determined_Rotational_correlation_time), *n_residues_* is the random number of residues, and 110 is the average molecular weight of an amino acid. The τ_c_ used to generate the relaxation rates was then selected from a uniform distribution between 0.5 and 1.5 times *τ_c_*_,*base*_. Because the majority of the reported order parameter (S^2^) values in the literature are closer to 1, the synthetic S^2^ values were calculated according to *S*^2^_*syn*_ = 1 – [*random*(0,1)]^2^, where *random*(0,1) is a random number uniformly distributed between 0 and 1, which has the effect of biasing *S*^2^_*syn*_ closer to 1. Fast ps-ns timescale local motions (τ_e_) were included and scaled based on S^2^. τ_e_ values were chosen from a uniform distribution between 0 ps and a maximum value that depended on S^2^, starting at 300 ps when S^2^ = 0.8 and continuously scaling up to 1000 ps as S^2^ reaches 0. An upper limit of 0.4 times τ_c_ was applied to τ_e_. These ceilings should ensure that τ_e_ is always in the fast-tumbling limit (i.e., ωτ_e_ << 1). Finally, µs-ms timescale conformational exchange (R_ex_) was added to 50% of the synthetic proteins. Within this random 50% of the synthetic proteins that had additional R_ex_, 10% - 25% of the residues were randomly selected to have exchange contributions between 1 and 25 s^-1^. Note, the single global (τ_c_) and three local (S^2^, τ_e_, and R_ex_) dynamics parameters correspond to “model 4” in the model-free formalism.

The synthetic dynamics parameters were then used to generate ^15^N R_1_ and R_2_ relaxation rates and {^1^H}-^15^N heteronuclear NOE values using the slow isotropic tumbling with spatially restricted faster local motions model within the model free formalism^12,13^

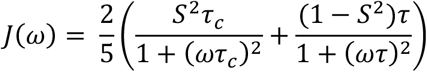

where τ^-1^ = τ_c_^-1^ + τ_e_^-1^. The relaxation inputs were calculated according to^14,46^

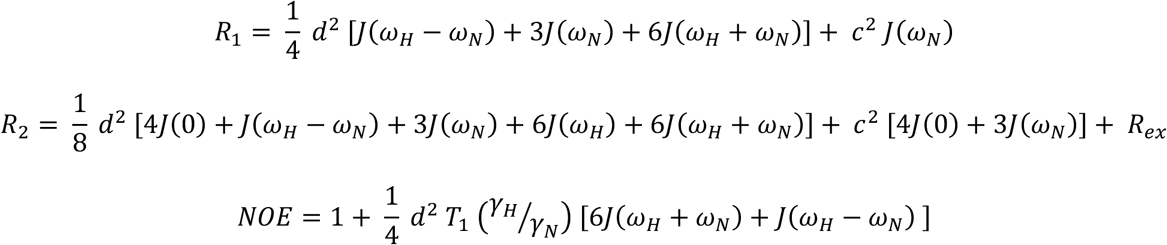

Where 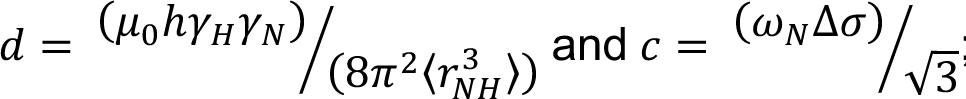; ω_H_ and ω_N_ are the Larmor frequencies for the ^1^H and ^15^N spins, respectively, µ_0_ is the permeability of free space, h is Planck’s constant, γ_H_ and γ_N_ are the gyromagnetic ratios for ^1^H and ^15^N nuclei, respectively, 〈*r_NH_*〉 is time-averaged length of the ^1^H-^15^N bond (1.02 Å), and Δσ is the chemical shift anisotropy for the ^15^N spin (−160 ppm). Random error, between 2% and 8% of the value, was added to the relaxation rates and NOE value. Features (i.e., R_1_ and R_2_ rates and NOE values), local labels (i.e., τ_e_, S^2^, and R_ex_), and global labels (i.e., τ_c_) for 10k proteins were saved in *tensorflow* record files (40 files total for the 400k synthetic proteins). Note, τ_c_ and τ_e_ times were saved in units of nanoseconds, units we found facilitated model training.

### Model Architecture and Hyperparameters

The multi-task neural network was implemented using the *Keras* API within *TensorFlow*^47^. To accommodate proteins of varying lengths, the model accepts variable-length input sequences composed of three relaxation features. A masking layer is applied immediately after the input to ensure padded sequence elements do not contribute to the network computations. The shared feature-extraction base of the model consists of two consecutive Bidirectional Long Short-Term Memory (BiLSTM) layers with 128 and 96 units, respectively, configured to return the full sequence outputs. This sequential representation is passed through a fully connected Dense layer consisting of 96 units with a Rectified Linear Unit (ReLU) activation function, followed by a Layer Normalization step to stabilize the learned features across the sequence. Following normalization, the architecture splits into two task-specific branches. The residue-specific dynamics (S^2^, τ_e_, and R_ex_) are predicted by the local branch using a time-distributed linear Dense layer consisting of 3 output units, mapping the normalized sequence directly to the three local biophysical parameters per residue. To predict the global rotational correlation time, the normalized sequence is passed through the global branch consisting of a multi-head self-attention layer to weight the relative importance of each residue. The attention-weighted sequence is then condensed using 1D global average pooling. This pooled, protein-level feature vector is processed by a Dense layer (64 units, ReLU activation) and regularized using a Dropout layer (rate = 0.3) to prevent overfitting before a final linear Dense layer with a single unit outputs the global τ_c_ prediction.

### Model training

To provide error estimation of the modeled dynamics parameters, an ensemble approach was utilized where ten models were generated using different training and validation sets from the synthetic training data. The final extracted dynamics parameters that SPINDLE reports are the mean and standard deviation from these ensembles. For each of the ten models, the list of the *tensorflow* record files is randomly shuffled, and the first 90% (i.e., 36) of the files (containing 360k synthetic proteins) were used for training, and the last 10% (i.e., 4) of the files (containing 40k synthetic proteins) were used for validation during training. Deep learning models were generated with the *tensorflow* v2.18 python package utilizing the *keras* v3.13 API. Optimization was performed within *tensorflow* using the ADAM optimizer^48^ and an initial learning rate of 5 x 10^-4^ which was lowered to 1 x 10^-6^ based on the loss calculated from the validation data between training epochs. Two loss functions were used to train the model: for the global branch (i.e., τ_c_), the mean absolute error (MAE) was calculated, whereas for the local branch (i.e., S^2^, τ_e_, and R_ex_), a weighted MAE was used. This weighted MAE scales the individual MAEs for S^2^, τ_e_, and R_ex_, since the local dynamics parameters have different ranges (S^2^: 0-1, τ_e_: 0-1 ns, and R_ex_: 1-25 s^-1^). To achieve the scaling, raw MAEs for S^2^, τ_e_, and R_ex_ were multiplied by 10, 10, and 0.4, respectively. Moreover, an additional penalty was added to the local error when the model predicted R_ex_ for a residue that does not have any exchange. Model training was performed using the high-throughput computing environment (*HTCondor*) on NMRBox^49^. Each model was trained with one GPU, requiring at least 8 Gb of RAM, and took 1-2 hrs.

### Post-hoc calibration

Initial testing of the models suggested that small post-hoc correction(s) may have been needed. To test the deep learning models for each magnetic field and determine whether any post-hoc corrections are needed, 10k new synthetic ‘proteins’ were generated using the same logic as described above. The resulting relaxation data were provided to the models, and the predicted dynamics parameters were compared to the ground truth values. Magnetic field specific linear corrections were applied τ_c_; no corrections were applied to the local parameters. The same validation/calibration data were also used to calibrate error multipliers to ensure that 95% of the ground truth values fall within the 95% confidence intervals of the predicted mean ± standard deviation of the SPINDLE ensemble predictions. Lastly, the validation/calibration data was used to define ambiguity thresholds. Here, ceilings were defined for the maximum error that can be tolerated before the prediction becomes misleading: 0.1, 250 ps, and 2.5 s^-1^ for S^2^, τ_e_, and R_ex_, respectively. Then, using the true absolute error between the ground truth values and the SPINDLE predictions, F_1_-scores (precision-recall curves) were determined; the ambiguity threshold was set as the value that maximized the F_1_-scores.

## Code availability

All the *python* code and synthetic training data generated during this work along with a stand-alone SPINDLE python script and Jupyter notebook of the SPINDLE script for Google Colab is freely available from GitHub (https://github.com/michaellatham77/SPINDLE) under the MIT General Public License Agreement.

## Acknowledgements

We thank Marella Canny (UMN) for critically reading this manuscript. This work was supported by the National Institutes of Health/National Institute for General Medical Studies (NIH/NIGMS) grant R35GM128906. This material is based upon work supported by the Air Force Office of Scientific Research under award number FA9550-25-C-B010 in the amount of $44,400.

## Author Contributions

M.P.L. conceived and designed the project. O.E.K. and M.P.L. developed the machine learning framework, performed data analysis, and prepared the manuscript.

