## Supplemental Figure for "SPINDLE: Unlocking protein dynamics from single-field NMR relaxation data using a deep learning ensemble"

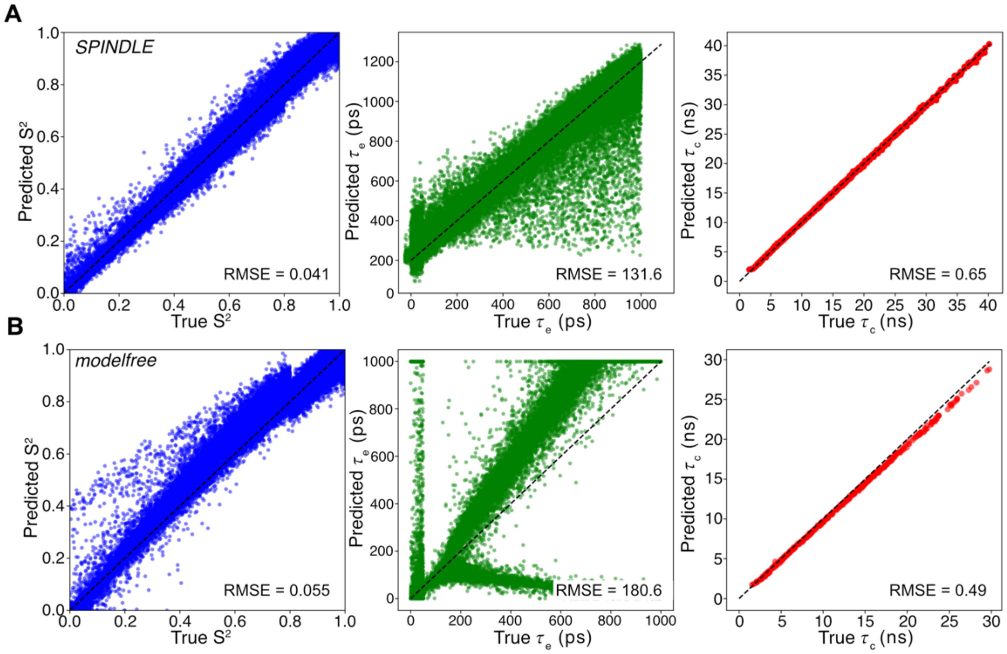


**Supplemental Figure 1. Comparison of SPINDLE predictions and conventional non-linear least squares analysis on synthetic 800 MHz model 2 dynamics data.** Comparison between (A) SPINDLE deep learning predictions and (B) traditional non-linear least-squares optimization (via *modelfree* v4) using synthetic model 2 (i.e., global $\tau_c$ and local $S^2$ and $\tau_e$) relaxation data generated for 1000 'proteins' at 800 MHz. *Left column*, predicted/calculated vs ground truth $S^2$; *middle column*, predicted/calculated vs ground truth $\tau_e$; *right* column, predicted/calculated vs ground truth $\tau_c$. In all plots, the black dashed line indicates the y=x line, and the root mean square error (RMSE) value comparing the predicted/calculated and ground truth values is reported in the lower-right corner of each plot.

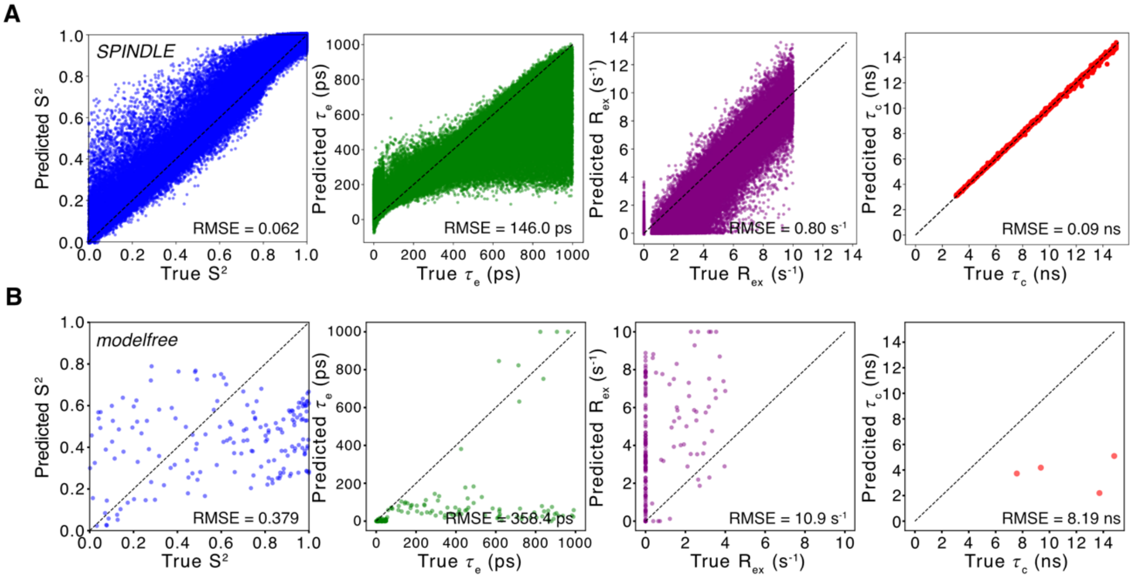


**Supplemental Figure 2. Comparison of SPINDLE predictions and conventional non-linear least squares analysis on synthetic 800 MHz model 4 dynamics data.** Comparison between (A) SPINDLE deep learning predictions and (B) traditional non-linear least-squares optimization (via *modelfree* v4) using synthetic model 4 (i.e., global $\tau_c$ and local $S^2$, $\tau_e$, and $R_{ex}$) relaxation data generated for 1000 'proteins' at 800 MHz. *First column*, predicted/calculated *vs* ground truth $S^2$; *second column*, predicted/calculated *vs* ground truth $\tau_e$; *third column*, predicted/calculated *vs* ground truth $R_{ex}$; and *forth column*, predicted/calculated *vs* ground truth $\tau_c$. In all plots, the black dashed line indicates the y=x line, and the root mean square error (RMSE) value comparing the predicted/calculated and ground truth values is reported in the lower-right corner of each plot.

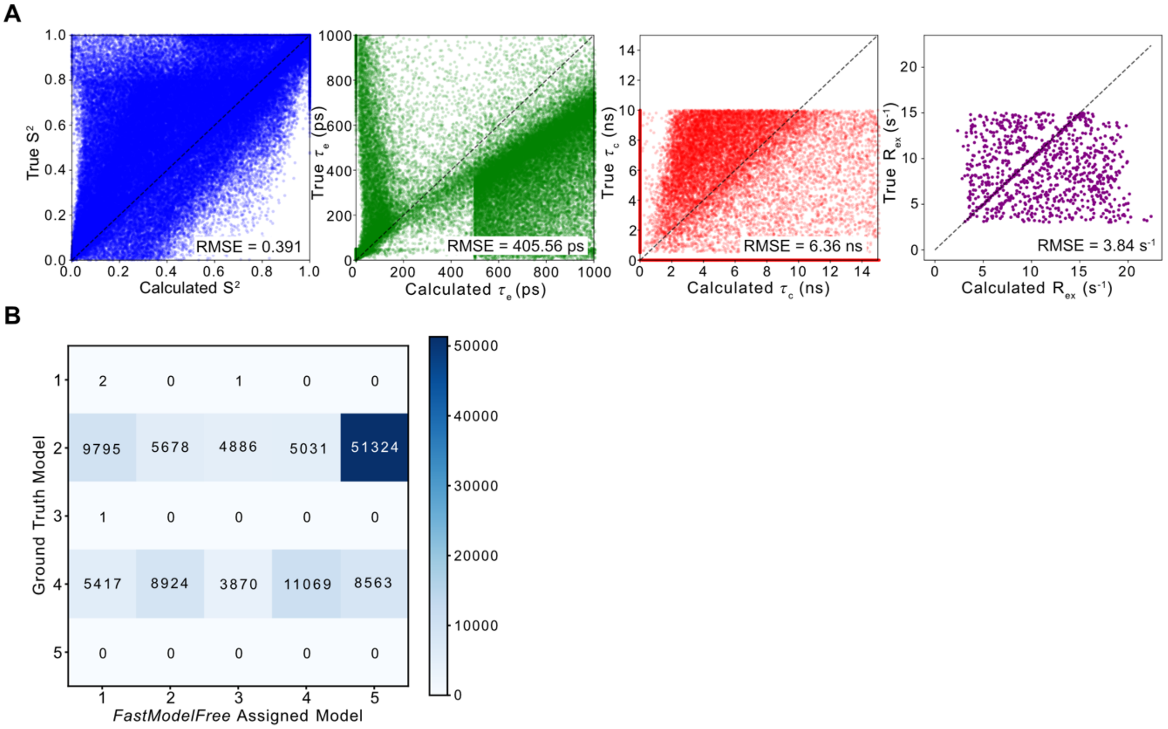


**Supplemental Figure 3. Evaluation of automated model selection on synthetic 800 MHz relaxation data.** (A) Comparison of ground truth *vs* non-linear least-squares calculated (via *FAST-modelfree and modelfree* v4) dynamics parameters from relaxation data generated for 1000 synthetic proteins at 800 MHz. *Left*, calculated *vs* ground truth $S^2$; *left middle*, calculated *vs* ground truth $\tau_e$; *middle right*, calculated *vs* ground truth $R_{ex}$; and *right*, calculated *vs* ground truth $\tau_c$. In all plots, the black dashed line indicates the y=x line, and the root mean square error (RMSE) value comparing the calculated and ground truth values is reported in the lower-right corner of each plot. (B) Heatmap comparing the actual model-free model selection of the synthetic ground truth data (y-axis) against the model selected by hypothesis testing within *FAST-modelfree* (x-axis). The numbers within each cell indicate the total number of assigned residues with the color scale corresponding to the residue count.

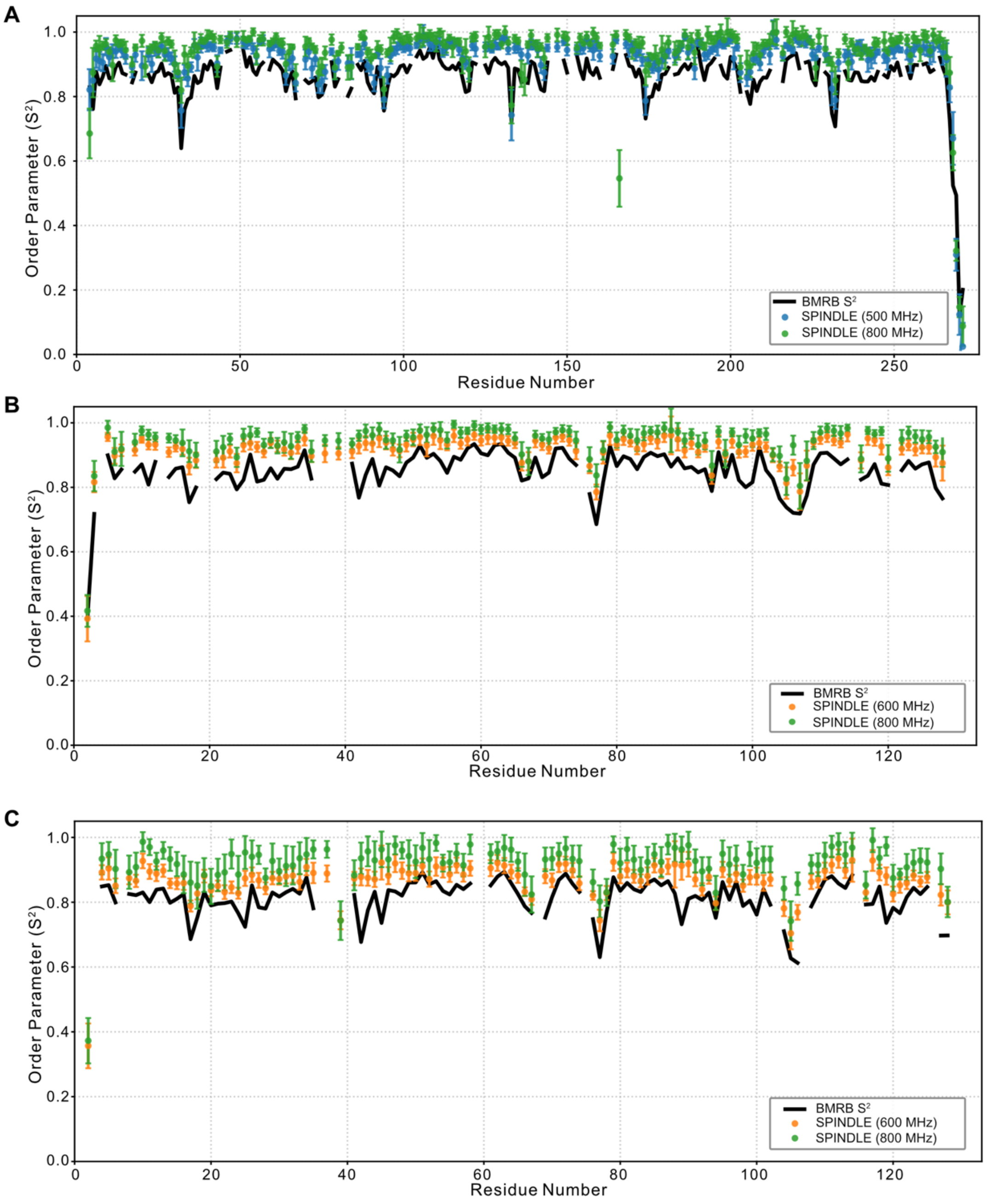


**Supplemental Figure 4. Comparison of SPINDLE-predicted $S^2$ with BMRB-deposited values at different field strengths and temperatures.** (A) Per-residue $S^2$ values for β-lactamase PSE-4 (BMRB ID: 6838) comparing BMRB-deposited values (solid black line) with SPINDLE predictions derived from single field relaxation data collected at 500 MHz (blue circles) and 800 MHz (green circles). (B,C) Per-residue $S^2$ values for azurin (BMRB ID: 6243) recorded at (B) 16 ºC and (C) 44 ºC comparing BMRB-deposited values (solid black line) with SPINDLE predictions derived from single field relaxation data collected at 600 MHz (orange circles) and 800 MHz (green circles). In all panels, the SPINDLE points and error bars represent the mean and standard deviation of the predictions across the 10-model ensemble.

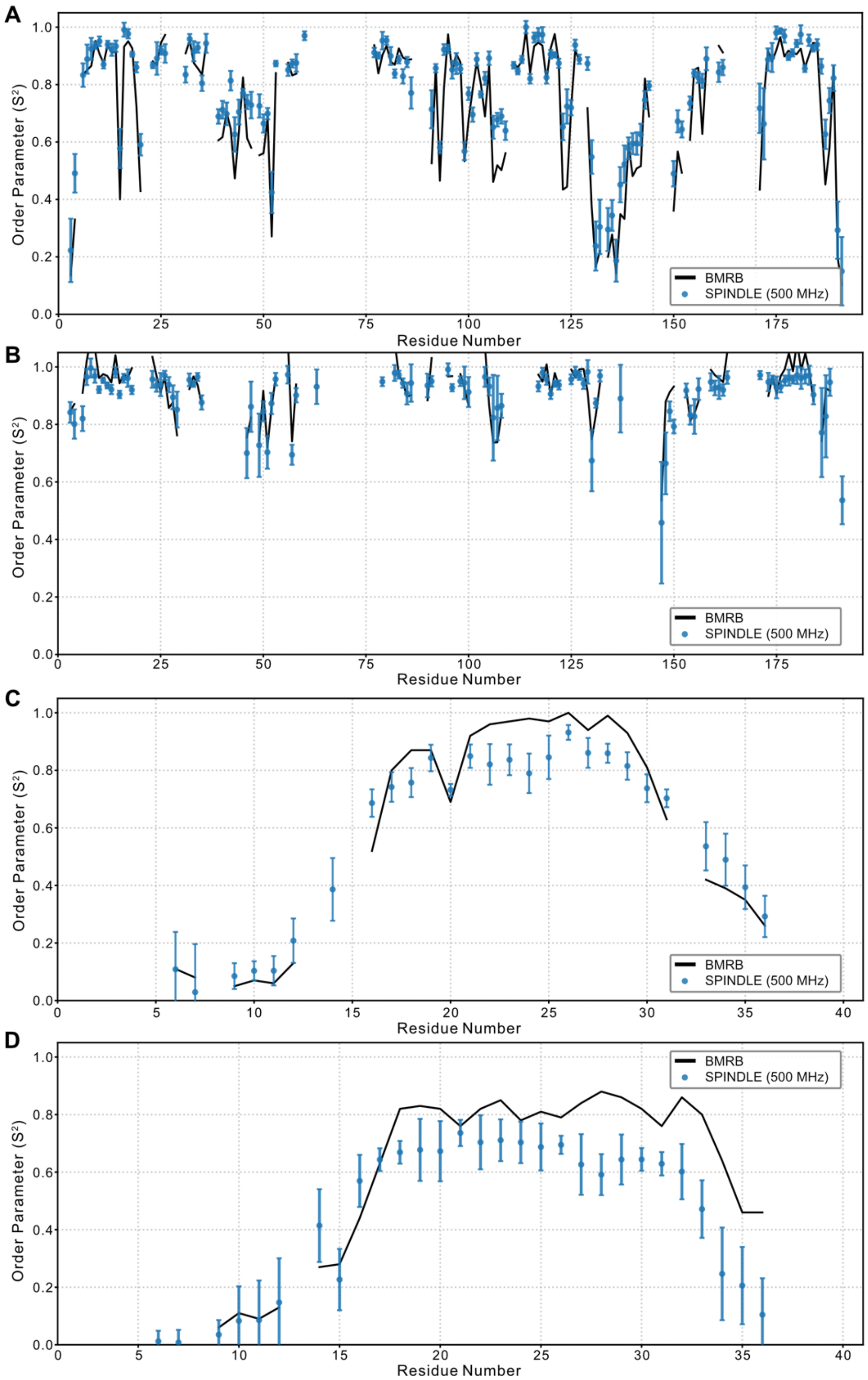


**Supplemental Figure 5. Comparison of SPINDLE-predicted $S^2$ with BMRB-deposited values under different pH conditions or for wildtype vs mutant.** (A,B) Per-residue $S^2$ values for recombinant human grown hormone (BMRB ID: 4689) at pH 2.7 (A) and pH 7.0 (B) comparing BMRB-deposited values (solid black line) with SPINDLE predictions derived from single field relaxation data collected at 500 MHz (blue circles). (C,D) Per-residue $S^2$ values for a wildtype (C;

BMRB ID: 5549) or mutant (D; BMRB ID: 5550) neuropeptide bound to dodecylphosphocholine micelles comparing BMRB-deposited values (solid black line) with SPINDLE predictions derived from single field relaxation data collected at 500 MHz (blue circles). In all panels, the SPINDLE points and error bars represent the mean and standard deviation of the predictions across the 10-model ensemble.

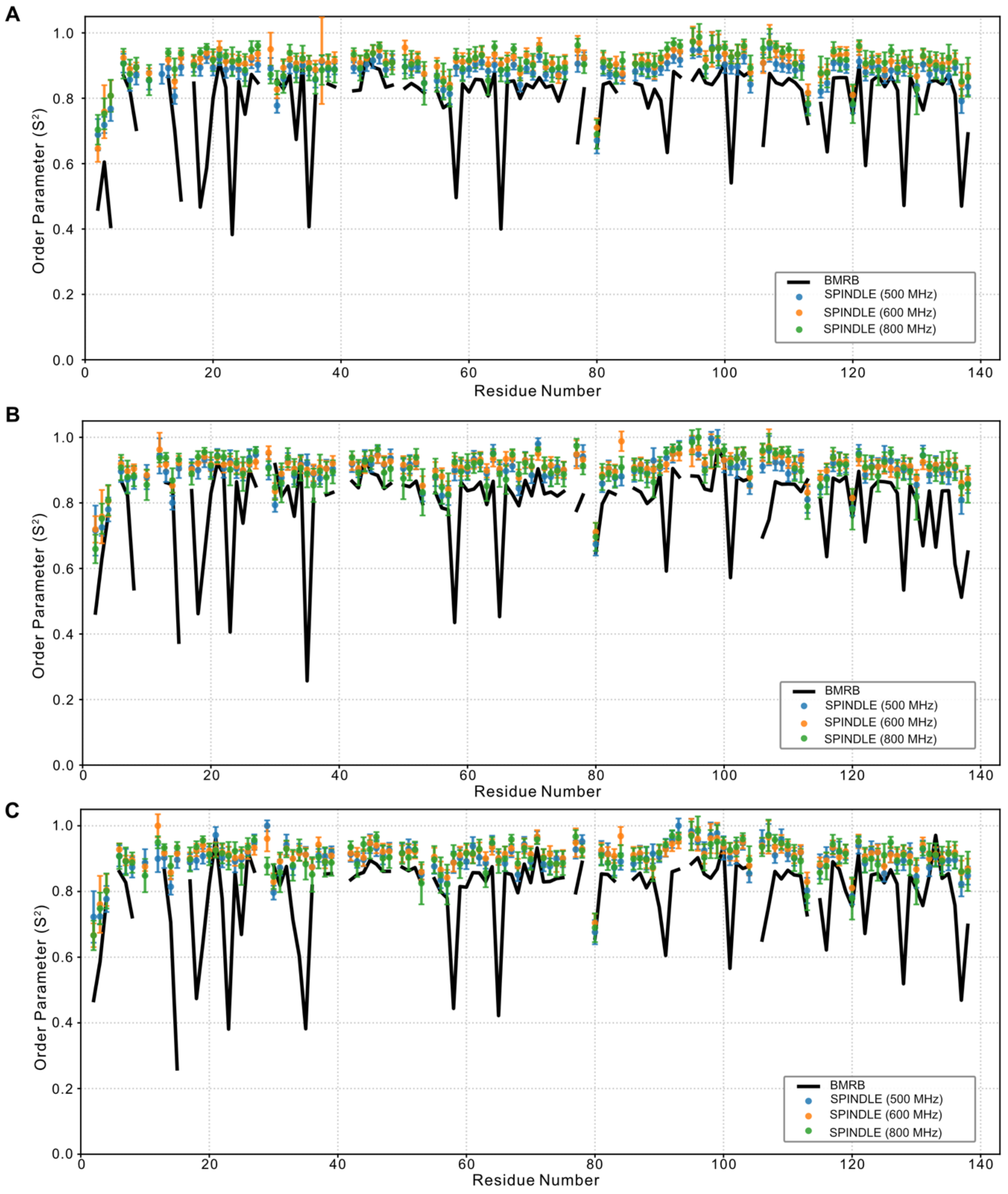


**Supplemental Figure 6. Comparison of SPINDLE-predicted $S^2$ with BMRB-deposited values for different galectin-3C carbohydrate complexes.** Per-residue $S^2$ values for galectin-3C bound to different carbohydrate substrates (A; BMRB ID: 50283), (B; BMRB ID: 50284), and (C; BMRB ID:50285) comparing BMRB-deposited values (solid black line) with SPINDLE predictions derived from relaxation data collected at 500 MHz (blue circles), 600 MHz (orange circles), and 800 MHz (green circles). In all panels, the SPINDLE points and error bars represent the mean and standard deviation of the predictions across the 10-model ensemble.

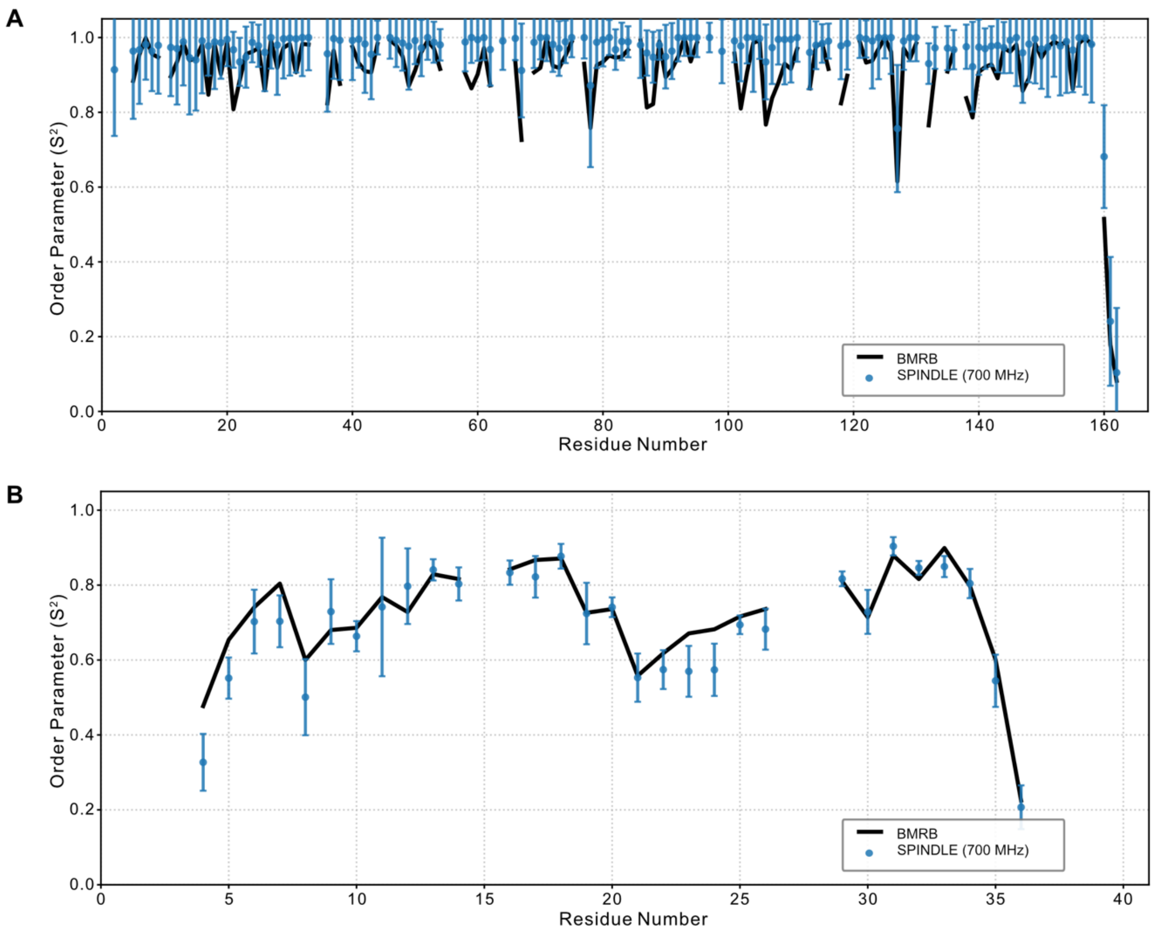


**Supplemental Figure 7. Comparison of SPINDLE-predicted $S^2$ with BMRB-deposited values.** (A) Per-residue $S^2$ values for the 110 kDa hexameric inorganic pyrophosphatase (BMRB ID: 50212) comparing BMRB-deposited values (solid black line) with SPINDLE predictions derived from relaxation data collected at 700 MHz (blue circles). (B) Per-residue $S^2$ values for the 34 residue di-GMP sequestering peptide (BMRB ID: 50001) comparing BMRB-deposited values (solid black line) with SPINDLE predictions derived from relaxation data collected at 700 MHz (blue circles). In all panels, the SPINDLE points and error bars represent the mean and standard deviation of the predictions across the 10-model ensemble.

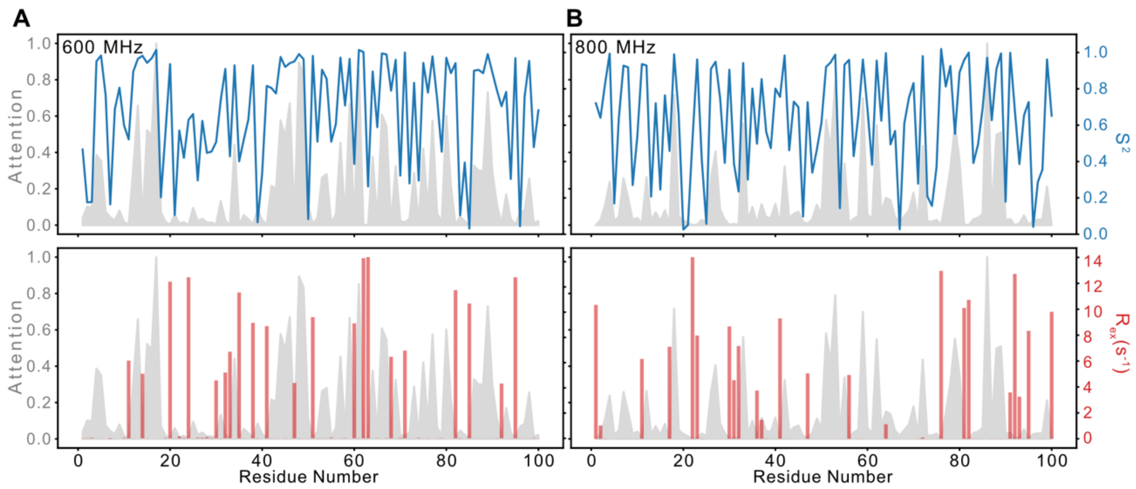


**Supplemental Figure 8. SPINDLE self-attention profiles highlight which residues are important for extracting dynamics parameters.** (A and B) Normalized per-residue attention weights extracted from the SPINDLE Multi-Head Self-Attention layer (grey shaded area, left y-axis) overlaid with predicted $S^2$ (blue lines, top subplot, right y-axis) and $R_{ex}$ (red bars, bottom subplot, right y-axis). Attention weights and predicted dynamics parameters were obtained from synthetic $^{15}N$ relaxation data calculated for 100-residue proteins with $\tau_c$ = 5.87 ns at 600 MHz (A) and $\tau_c$ = 6.74 ns at 800 MHz (B).
